# Dominant baboons experience more interrupted and less rest at night

**DOI:** 10.1101/2025.01.06.629345

**Authors:** Marco Fele, Charlotte Christensen, Anna M. Bracken, M. Justin O’Riain, Miguel Lurgi, Marina Papadopoulou, Ines Fürtbauer, Andrew J. King

## Abstract

The amount and quality of sleep individuals get can impact various aspects of human and non-human animal health, ultimately affecting fitness. For wild animals that sleep in groups, individuals may disturb one another, influencing sleep quality and quantity, but this aspect of social sleep has been understudied due to methodological challenges. Here, using nighttime rest (absence of bodily movements) as a proxy for sleep, we test the hypothesis that individual’s social dominance can affect sleep opportunities by studying a troop of wild chacma baboons (*Papio ursinus*), a species with a strong hierarchical social structure. First, we show that the troop’s night-time rest (determined by 40Hz acceleration data) is highly synchronised. Next, we link night-time rest dynamics to daytime spatial networks and dominance hierarchy (from 1Hz GPS data and direct observations). We show that baboon rest synchrony is higher between similarly ranked individuals, and unexpectedly, more dominant baboons experience less and lower-quality rest. We propose that this hierarchy effect is explained by higher-ranked baboons resting closer to more group members, which also leads them exerting greater influence on each other’s night-time behaviour compared to lower-ranked individuals. Our study provides the first empirical evidence for the impact of social hierarchies on sleep in a wild primate, suggesting that dominance status may impose trade-offs between social rank and the quality and quantity of sleep.

## MAIN TEXT

Sleep is a fundamental biological process (Xie et al. 2013; Klinzing et al. 2019). For group living animals that sleep together – such as humans (*Homo sapiens*) – communal sleep promotes safety (Samson et al. 2017) but the restlessness of nearby individuals can disturb sleep (Crittenden et al. 2018). Similar disruptions occur in other social primates like baboons (*Papio ursinus*) (Loftus et al. 2022) and macaques (*Macaca fuscata yakui*) (Mochida and Nishikawa 2014), leading to the synchronisation of sleep/wake behaviour among group members. However, the disruptive effects of others’ activity may not be equal for all group members, especially in groups with social hierarchies where more dominant baboons may monopolise the best sleeping positions (Smeltzer et al. 2022). The effect of specific social structures like dominance hierarchies on sleep is therefore an important but understudied aspect of social sleep, largely because of the challenges associated with collecting and combining detailed daytime and night-time behavioural data for known individuals in natural settings (King et al. 2018; Smeltzer et al. 2022; Chakravarty et al. 2024).

Here, we study the social dynamics of rest for individually identifiable members of a wild chacma baboon (*Papio urinus*) troop in the Cape Peninsula, South Africa. Pairing information on daytime spatial networks and social hierarchies constructed from direct observations and GPS data, with analyses of night-time activity from accelerometer data, we test the hypothesis that social status impacts rest duration and quality in social animals. We predicted dominant individuals benefit from higher quality sleeping positions and, in turn, better sleep (Smeltzer et al. 2022). Baboons are an ideal model system in which to explore the potential effect of a group’s social structure on sleep (Loftus et al. 2022; Chakravarty et al. 2024). Chacma baboon groups comprise a complex social system with a linear dominance hierarchy (Silk et al. 1999; Cheney and Seyfarth 2008; Silk et al. 2010; Cheney et al. 2016). The dominance hierarchy is stable and mediates association patterns among group members, with higher ranked baboons occupying central positions in social networks (King, Clark, et al. 2011; Fele et al. under revision). Furthermore, the importance of higher-ranked individuals is critical to group-level processes during the daytime (e.g. group decision-making and coordination: King et al. 2008; Stueckle and Zinner 2008; Stueckle and Zinner 2008; Kaplan et al. 2011; King, Sueur, et al. 2011; Sueur 2011; Bracken, Christensen, O’Riain, Fürtbauer, et al. 2022).

### Baboons synchronise rest

We use rest (absence of physical activity) as a proxy for sleep (see methods). Individuals spent 9.0±0.19 hours resting and 1.37±0.19 hours active each night (mean ± SD) (Figure 1A) and were active earlier in the morning and rested later in the evening with increasing day length, resulting in less hours of rest (Generalised Linear Mixed Model [GLMM], beta: −0.01, p < 0.001; Fig S1). For approximately one third of the night, all individuals were sleeping simultaneously (2.78 ± 0.69 hours per night); much more time than would be expected if all individuals behave independently (Chi-square goodness of fit: p < 0.001; Fig 1B). By modelling the resting and active states of individuals as a Markov chain (Fig 1C), we tested if (i) rest quantity, 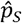 (the overall probability of resting during the night), (ii) sleep quality, 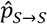 (i.e., the probability of resting in successive timesteps), and (iii) wakefulness, 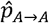, (i.e. the probability of being active in successive timesteps) were predicted by the number of other resting individuals. We found that with more resting individuals, individuals’ rest quality 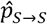 increases (GLMM: beta: 0.20, p < 0.001) and wakefulness 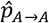 decreases (GLMM: beta: −0.06, p < 0.001) resulting in an overall increase in rest quantity 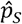 (Fig 1D) when baboon rest is synchronised. In our analysis, we also controlled for the effects of night-time rainfall, temperature, and moon illumination, and the temporal autocorrelation in rest across nights (see Table S1 and S2 for full model results).

**Figure 1.**
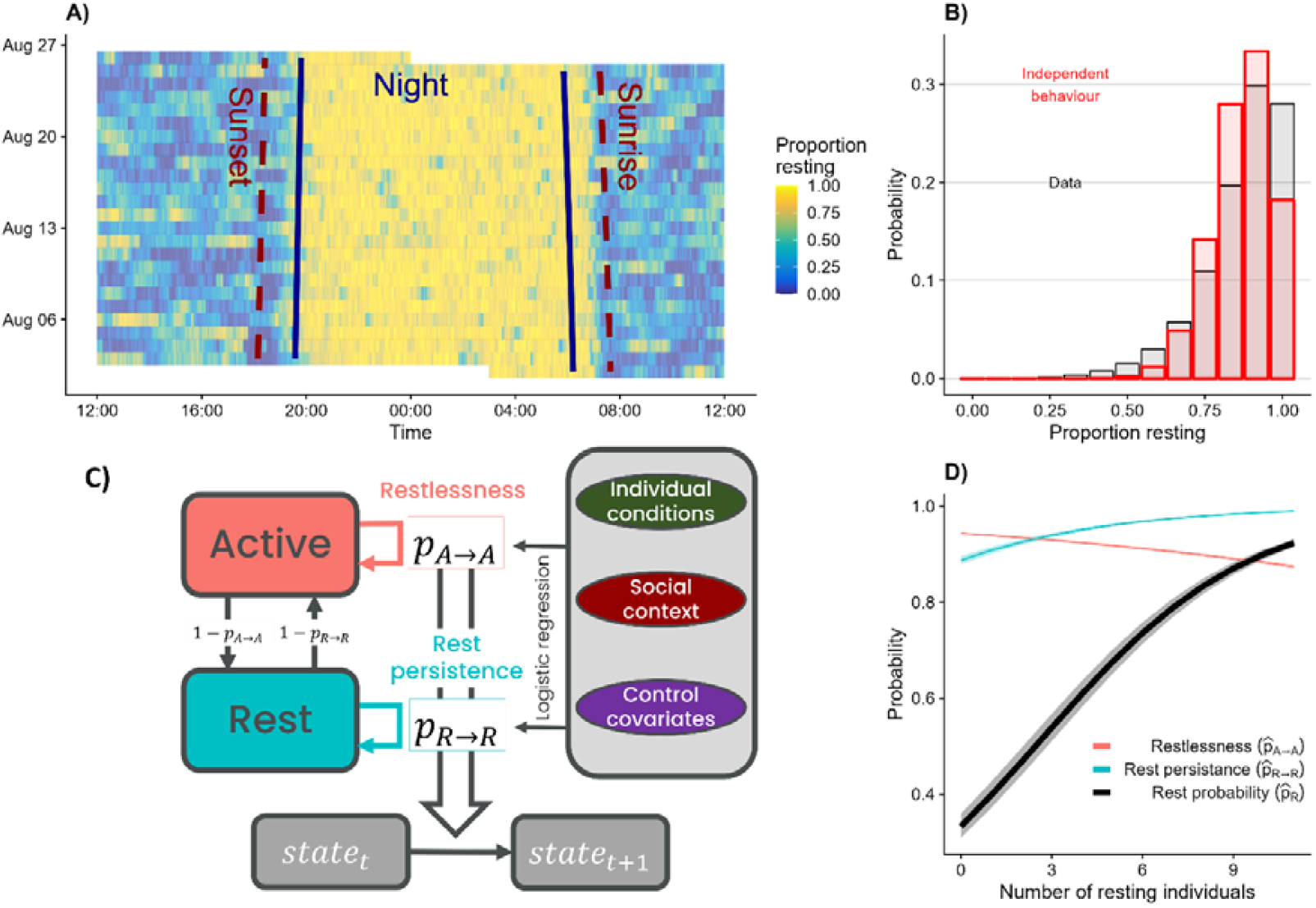
Baboon rest dynamics. (A) Proportion of baboons wearing tracking collars that are at rest (mean VeDBA <0.04; see methods) over days (y axis) and time (x axis). Night darkness, sunrise, and sunset are indicated by vertical lines. (B) Observed probability a given proportion of baboons wearing tracking collars are recorded as resting (top) compared to the probability expected for a null model of independent rest (see methods). (C) Using the probability of baboons switching between active and rest states we model rest quality, (i.e., the probability of resting in successive timesteps), and wakefulness,, (i.e., the probability of being active in successive timesteps) giving us overall rest quantity, (the overall probability of resting during the night). In analyses, we control for individual, social and environmental factors. (D) Applying our framework in C we find that when more baboons are resting, the rest quality (staying inactive) goes up, wakefulness (staying awake) goes down, and rest quantity increases. The number of resting individuals (x-axis) excludes the focal individual.

### Dominant baboons have less and lower quality rest

Baboons of higher dominance rank had lower rest quality (GLMM: beta: −3.52, p < 0.001) and higher wakefulness (GLMM: beta: 2.32, p < 0.001), resulting in them getting overall less rest (Fig 2A, Fig S2), from ~8.5 hours to ~6 hours (Fig 3S, 4S). To understand this hierarchy effect on rest, we considered *how* baboons may influence each other’s rest. Given our synchrony result (above) we tested if dominance mediated individuals’ response to their social environment and tested for an interaction between dominance and the number of other resting individuals. We found a positive interaction for rest quality (GLMM; beta: 0.04, p < 0.001), and a negative interaction for wakefulness (GLMM; beta: −0.04, p < 0.001), suggesting stronger rest synchrony effects among dominant individuals (Fig 2B). Visualising the observed synchrony in state for pairs of baboons (Fig S5) suggests that individuals of similar dominance have a higher probability of being in the same state, which is also endorsed by our model (interaction : 4.97, p-value < 0.001; interaction : −3.11 p-value < 0.001, see Fig S6, S7, S8 for model predictions). We therefore performed a cross-correlation analysis (e.g. Nagy et al. 2010; Fürtbauer et al. 2020; Georgopoulou et al. 2022) of the behavioural time series (state) for each pair of individuals (see methods). We found that the maximum cross-correlation was higher between dominant individuals (GLMM: beta: 0.26, lower credible interval (CI) = 0.02, upper CI = 0.48, Fig 2D), with more dominant individuals being on average the “leaders” in the behavioural coupling (Fig 2E). This translates into dominant individuals having greater influence on others’ rest state (positive shifts of peak-cross correlation) (GLMM, beta: 0.44, p = 0.02 for lead/follow response; Fig 2F). Furthermore, females influence others’ state more than males (GLMM, beta: −0.42, p = 0.01), but note that our sample size for males is n = 2.

**Figure 2.**
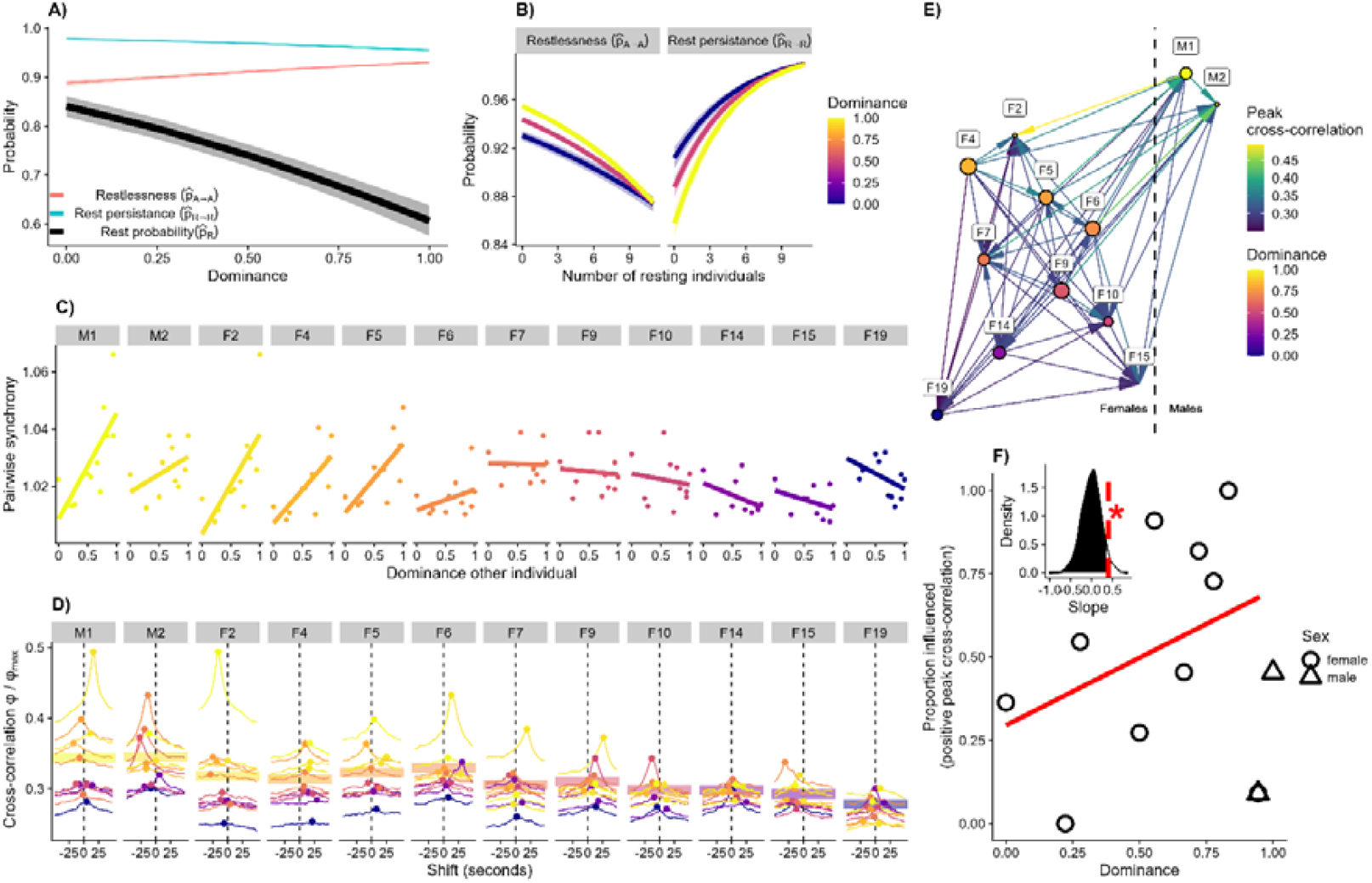
Dominance effects on baboons’ rest. (A) Higher ranked baboons have poorer rest quality, 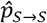 (i.e., the probability of resting in successive timesteps), are more likely to stay awake 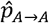, (i.e., the probability of being active in successive timesteps) and as a result have lower rest quantity, 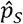 (the overall probability of resting during the night). (B) With more baboons resting (x axis) there is a higher probability of staying resting 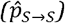 and lower probability of staying awake 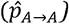 but these effects are moderated by dominance. (C) Pairwise synchrony is state (rest, awake) for each baboon as a function of the dominance of a paired individual. Baboons are ordered from left to right according to dominance. Raw data and a simple linear model are shown to illustrate the direction of the relationship – showing that similarly ranked baboons are synchronised. (D) Cross-correlation of a baboon state (rest, awake) with the state of others (ordered left to right by dominance). The strength of the cross-correlation (y axis) shows how similar the two baboons states are. The time shift (x axis) indicates whether other baboons change state before or after the focal baboon. Horizontal lines indicate average individual cross-correlation strength (E) Dominant baboons have greater influence on others rest; this figure is a network visualisation of the effect shown in D, where dominant baboons (left hand side in D) tend to have higher correlation with other high-ranked baboons (and tend to lead transitions). F) Correlation between individual dominance and the proportion of influenced baboons. Inset shows the distribution of dominance effects under a null model and the observed significant relationship (dotted line). See methods for more details.

**Figure 3.**
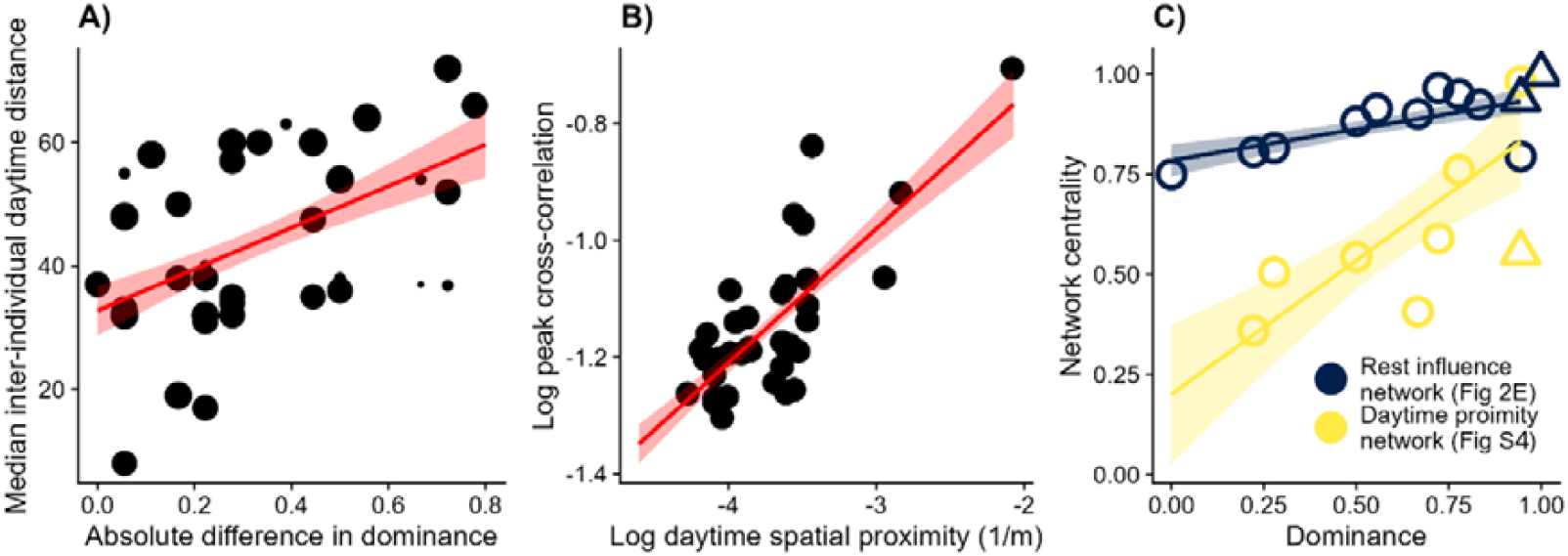
Proximity networks and rest influence. (A) Baboons of similar rank (x axis) spend more time closer to each other (y axis). Raw data are shown, with point size indicating sample size for estimating median daytime pairwise distance and a linear relationship is fitted for illustration. (B) Baboons that are closer to each other during the daytime (x axis) have higher correlation in their transition between rest and active states at night (y axis). (C) Higher ranked baboons (x axis) have greater eigenvector network centrality in daytime proximity networks and night-time rest influence networks (y axis).

### Why do dominant baboons have worse rest?

Together, our results show that more dominant baboons exert greater influence on each other’s night-time behaviour compared to lower-ranked individuals. We therefore tested whether a simple mechanism based on spatial proximity could explain these dynamics. We found that baboons of similar dominance rank are closer to each other during the day (GLMM, beta: 0.37, lower CI = 0.13, upper CI = 0.62, Fig 3A), and that daytime spatial proximity between baboons predicted rest influence at night (GLMM, beta: 0.83, lower CI = 0.61, upper CI = 1.03; Fig 3B). Furthermore, dominant baboons are more central (higher eigenvector centrality) in both daytime proximity networks (Fig S9) and the predicted rest influence network (Fig 2E) (GLMM proximity, beta: 0.67, p = 0.002; GLMM influence, beta: 0.15, p = 0.04). Therefore, we propose that dominant baboons have more, and closer neighbours at night-time, and so they end up waking, and being woken by other baboons more frequently (Fig 2C).

### The consequences of social structure on sleep

Sleep synchrony in social animals is a well-known phenomenon. For example, in colonies of seabirds, individuals sleep when their neighbours are also sleeping, resulting in waves of sleep propagating (Beauchamp 2008; Beauchamp 2011). Recent technological and theoretical advances have allowed for a better understanding of sleep as a collective phenomenon (Loftus et al. 2022; Chakravarty et al. 2024). Individuals influence each other’s night-time behaviour (Alberts 2019), and tend to synchronise their sleep with specific resting partners (Sotelo et al. 2024), especially when these partners are close kin or affiliates (Takahashi 1997; Lock and Anderson 2013; Brividoro et al. 2021). We observed similar rest synchronization in our study group. While we did not measure continuous environmental variation overnight that might simultaneously cause waking (though we controlled for night-level environmental effects; see Supplemental Results), our cross-correlation analysis supports direct social influence on rest, indicating individuals affect each other’s rest behaviour.

Our study provides the first evidence that dominance hierarchies influence rest dynamics in a wild animal. Data indicate that dominant individuals might occupy more central sleeping positions or sleep in bigger subgroups with more nearby neighbours, as also observed in sleeping lemurs (Radespiel et al. 1998; Lutermann et al. 2010). These positions reflect daytime associations, as seen in howler monkeys (*Alouatta caraya*) (Brividoro et al. 2021). Such association patterns lead to higher-ranked individuals exerting greater influence on each other’s night-time behaviour compared to lower-ranked individuals, resulting in more dominant baboons experiencing less and lower quality rest. Despite the recognised importance of social hierarchies and structure on animal lives during the daytime (King et al. 2008; King, Clark, et al. 2011; Majolo et al. 2012; Holekamp and Strauss 2016; Bracken, Christensen, O’Riain, Fürtbauer, et al. 2022) to our knowledge, only one study has investigated dominance rank effects on sleep, studying groups of freely moving mice (Karamihalev et al. 2019). In this study, authors found that socially dominant individuals had overall reduced slow-wave activity and more fragmented sleep but were unable to identify a cause for these differences. Future work can now investigate whether social disruption—a mechanism we propose—is widespread and explore its potential consequences.

Our data suggest that dominant individuals may experience rest costs due to their social integration, though they might also gain benefits. For animals sleeping in the open, dominant individuals are predicted to compete for central positions (Smeltzer et al. 2022) because they may confer lower predation risk (Yagi and Hasegawa 2011; Picman et al. 2022), thermal benefits (Takahashi 1997) or decrease vulnerability to insect bites (Mooring and Hart 1992). Even if dominants do not compete for central positions they may ‘end up’ at the centre of sleeping aggregations simply because of their many and strong social bonds (Farine et al. 2017; Fele et al. in revision), a result of spatial and social processes being intertwined (Webber et al. 2023; Albery et al. 2024). Further work should now focus on extending and testing the mechanisms we have uncovered here to other social sleeping species with strong dominance hierarchies.

Because dominant baboons experience less and lower quality rest than subordinates, social rank may result in unequal rest-related costs and benefits. Reduced and interrupted sleep can influence various physiological and cognitive functions (Xie et al. 2013; Klinzing et al. 2019), impacting health and fitness (Rechtschaffen 1998). Disrupted sleep can impair decision-making (Wild et al. 2018), weaken immune function (Besedovsky et al. 2019), and even result in death (Rechtschaffen et al. 1983). Whether more dominant baboons in fact incur costs because of “social sleep disruption” remains to be tested. Future work can now do this by, for example, linking variation in sleep quality and quantity to other behavioural and physiological measures (Meerlo et al. 1997; Fürtbauer et al. 2020; Christensen et al. 2022; Morgan et al. 2023). However, dominant individuals may actually require less rest because of some other correlated trait (Noser et al. 2003; Shuboni-Mulligan et al. 2021), or individuals may not be rest limited, making differences in loss or disruption of rest across individuals negligable.

The high levels of rest synchrony we have found suggests that individuals are not taking turns at resting to be prepared for external dangers as predicted by the “sentinel hypothesis” (Burger et al. 2020; McKinnon et al. 2023). Night-time leopard attacks are one of the main causes of death for baboons (Bidner et al. 2018), and so one would expect strategies to alert the group to risk (though this would be historical since our troop lives in a predator-free region of South Africa). Because dominant baboons (which we assume to be more central and to have more neighbours) are more likely to cause waking of others when they wake, it could be that they behave as ‘indirect’ sentinels. In fact, aggregation patterns of sleepers could be fine-tuned to propagate information – thus successfully identifying risks but minimising disruptions (Fahimipour et al. 2023; Gómez-Nava et al. 2023; McCormick et al. 2024), and testing this idea would be an interesting direction for future work.

## Conclusions

Our observations of wild baboons indicate that social hierarchies impact rest dynamics. Higher-ranked baboons – being more connected in spatial networks exert greater influence on each other’s night-time behaviour compared to lower-ranked individuals, resulting in less and more interrupted rest. Given that higher dominance in social animals is linked to better health (Sapolsky 2004; Sapolsky 2005) and longer lifespans (Campos et al. 2020; Snyder-Mackler et al. 2020) future work must now investigate if and how dominance status may impose costs as a consequence of bad sleep.

## Acknowledgments

We thank the Baboon Technical Team, Cape Nature and SANParks for authorization to conduct our research. We also thank Human Wildlife Solutions, field assistants Lucy Robertson, Charlotte Solman and Francesca Marshall-Stochmal, vets Dorothy Breed and Gary Buhrman, Esme Beamish and Layla King for their help and support. We thank Lucia Pedrazzi and Gui Araujo for their helpful discussion. We thank the South African Environmental Observation Network for weather data.

## Author contributions

M.F.: data curation, formal analysis, investigation, methodology, visualization, writing—original draft, writing—review and editing; A.M.B. data curation, investigation, methodology; C.C..: data curation, investigation, methodology; M.L.: resources, supervision, writing—review and editing; M.P.: data curation, formal analysis, visualization; writing—review and editing; M.J.O.: project administration, resources; I.F.: funding acquisition, project administration, resources, supervision, writing—review and editing. A.J.K.: funding acquisition, project administration, resources, supervision, writing—original draft, writing—review and editing.

## METHODS

### Study Site and Troop

We studied the ‘Da Gama’ baboon troop in Da Gama Park, Cape Town, South Africa (−34.15562° N, 18.39858° E). The troop consisted of approximately 50 individuals (2 adult males and 19 adult females). Baboon groups on the Cape Peninsula range near to and within urban areas of the City of Cape Town, and 13% of this study troop’s home range is classified as urban space (Bracken, Christensen, O’Riain, Fehlmann, et al. 2022). During the period for which we have full night-time data (see below), the troop slept on an apartment block in the urban space except for three nights when they slept in natural space. Rest was comparable in all qualitative comparisons we made between the natural and urban rest sites (Fig S10). Field observations also indicate the troop most probably splitting in subgroups for all but 9 nights, but we do not control for this in the analysis because we do not possess GPS night data and cannot determine with certainty subgroup composition.

### Tracking Collars

Most adults in the study troop (16/22) were fitted with tracking collars (for full details see Christensen et al. 2023 and Bracken et al. 2021). Use of collars and daily follows were approved by Swansea University’s Ethics Committee (IP-1315-5) and local authorities (Cape Nature, permit number: CN44-59-6527; SANparks, permit number: CRC/2018-2019/008-2018/V1). Collars contained tri-axial accelerometer data recorded at 40 Hz for 24hrs a day, and GPS tags (GiPSy 5 tags, TechnoSmArt, Italy) recording baboon position between 08.00h and 20.00h local time (UTC+2) at 1Hz. GPS positional accuracy was within 5m (and often much less than this) and erroneous fixes (average of 0.01% of GPS points per collar) were removed and interpolated, as described in Bracken et al. (2021). Because of collar failures, we have full accelerometery data for 12 (2/2 males, 10/19 females) for 25 astronomical nights between 2^nd^ August 2018 and 27^th^ of August 2018, and daytime GPS between 30/7/2018 and 15/9/2018 for 13 individuals (2/2 males, 11/19 females) (Table S3 and table S4). This resulted in both accelerometer and GPS data for 9 (2/2 males, 7/19 females) individuals.

### Daytime Spatial Networks and Dominance Rank

GPS data was used to study daytime proximity for the individuals for which we have spatial data. We constructed a network of inter-individual distances (or pairwise spatial associations). We represent every individual as a node, and each edge is the median pairwise individual distance. We use median pairwise distances for our network edges because we have data for collared individuals (Fig S11), with no need to control for sampling effort. Behavioural data on dominance interactions was collected *ad libitum* during direct observation over 78 days. Dominance rank was determined based on the outcome of directly observed dyadic agonistic interactions and then standardised between 0 (lowest) and 1 (highest).

### Night-time Rest

Astronomical night is defined as when the geometric centre of the sun is 18 degrees below the horizon, occurring approximately between ~20:00 and ~06:00 in our study system (Table S5) We obtained this data with the R package “suncalc” (Thieurmel and Elmarhraoui 2017). Accelerometer data was used to study rest. For every second in our accelerometer data, we calculate the average vector of dynamic body acceleration (VeDBA) (for details see Christensen at al. 2023) and consider an individual at complete rest when mean VeDBA are below 0.04, and active otherwise. VeDBA this low is well within the range of “rest” behaviour previously validated by comparing accelerometers output and video footage (Christensen et al. 2023). Proxies of sleep similar to ours have already been used (Gravett et al. 2017), and our sensitivity analyses (with threshold 0.03 and 0.05) suggest that our results are robust to different thresholding values (Fig S12, see Table S6, S7, S8, S9 for model results for sensitivity analysis).

### Rest Synchrony

To investigate group level rest synchrony, we compared the observed distribution of the probability of finding *n* individuals simultaneously resting with a null distribution in which individuals behave independently of each other. To calculate this, we first found for every individual *i* the observed probability of resting *p*_*i,S*_ as the total proportion of time spent resting. We then calculated the probability of a unique combination of resting and active individuals by multiplying together the probabilities of resting *p*_*i,S*_ for the resting individuals with the probabilities of being active 1-*p*_*i,S*_ for the active individuals. We find the null probability of *n* individuals resting by summing all the probabilities of having a group combination where *n* individuals are resting. We used a similar approach to investigate rest synchrony between pairs of individuals. We defined the null expectation of pairwise synchrony for individual *i* and *j* as *p*_*i,S*_ * *p*_*jS*_ + (1 − *p*_*i,S*_) *(1 − *p*_*j,S*_), which is the probability both are resting plus the probability both are active in case they behaved independently. We quantified the observed pairwise synchrony by calculating the proportion of time two individuals are in the same state in the data and divide this with the null expectation. This metric therefore provides information on how likely it is for two individuals to be in the same state, compared to what we would expect if their behaviour was independent.

### Rest Quantity, Rest Quality, and Wakefulness

We predicted: (i) rest quantity, 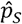 (the overall probability of resting during the night), (ii) rest quality, 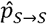 (i.e., the probability of resting in successive timesteps), and (iii) wakefulness, 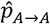, (i.e., the probability of being active in successive timesteps). We did this by considering the time series of “resting” and “active” of each individual as an inhomogeneous Markov chain (Fig 1C) with two states (Kim et al. 2009). “Inhomogeneous” refers to the state transition probabilities changing depending on social and environmental variables that vary in time (described below).

To find the rest quantity 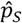, we calculated 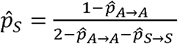, assuming the Markov chain has reached steady state (for details on model derivation, assumptions, and calculation of confidence intervals for 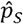 see supplementary material). To find the predicted transition probabilities 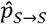 and 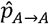 of the Markov chain, depending on the current state *s*_*t*_ at time *t*, we fitted two logistic regression models using the R package glmTMB (Brooks et al. 2017). Specifically, we fitted one logistic regression for when an individual is resting, and one when it is active (Fig 1B). The response variable for both logistic regressions is whether an individual is resting (1) or active (0) in the next time ste *S* _*t*+_l. To control for individual-level effects (non-independence of rest on successive nights), we included the observed rest probability during the previous night as a continuous variable and individual identity and night number as a random effect. To test how dominance impacts rest, we included standardised dominance rank as a continuous variable. To test whether the state of others changes the probability of transitioning into resting or active, we added the number of resting group members as an explanatory variable. We also included the average dominance of the resting individuals in interaction with individual dominance, to test if an individual is more or less likely to transition when the resting individuals are on average of higher, lower, or similar dominance. We also included environmental factors as controls (Fig 1B). We fitted rain conditions (yes or no; because rest may be disturbed or activity reduced when it rains), the average temperature (°C; because activity may be reduced on colder nights), moon illumination (fraction of the moon that is visible; because activity may increase on brighter nights), night duration (seconds, because baboons may rest proportionally less on longer nights). We gathered weather and temperature data at one hour resolution from the Cape Town weather stations of the South African Weather Service. We obtained moon illumination with the R package “suncalc” (Thieurmel and Elmarhraoui 2017), with one indicating full moon and zero indicating new moon.

### Social disruption of rest

We performed a cross-correlation analysis (Nagy et al. 2010; Fürtbauer et al. 2020; Georgopoulou et al. 2022) of the behavioural time series for each pair of individuals state at different temporal shifts. To do so, we temporally shifted the start of every focal individual’s timeseries and found the pairwise cross correlation with every other individual’s time series for different time lags (considering shifts spanning from −250 seconds to shift +250 seconds, see Supplemental Methods). The correlation coefficient for a binomial timeseries is Ψ/Ψ_*max*_ (Davenport and El-Sanhurry 1991), where Ψ is the covariance between the timeseries and Ψ_*max*_ is the maximum possible covariance. A peak in the cross correlation indicates the time lag for which the behaviour of the focal individual preferentially predicts the behaviour of the other. If the peak is positive, the focal individual is a “leader” and influences the other. If the peak is negative, the focal individual is a “follower” and is influenced by the other. Temporal autocorrelation of the timeseries (which is accounted in the previous model by using a Markov chain) does not bias the calculation of the cross correlation coefficient Ψ/Ψ_*max*_ (Dean and Dunsmuir 2016).

We constructed an influence network for rest disruption across our dataset, where every individual is represented as a node, and directed edges go from the influencer to the influenced individuals, with an edge value equal to the peak cross correlation. To test whether dominance predicts influence, we performed nodal regression (Hart et al. 2023) by calculating the proportion of interactions in which the focal individual influences others (was a leader, see above). We then fitted a logistic regression with individual dominance and sex as predictors, and the proportion of influenced individuals as response. To test whether the effect of dominance is significantly bigger than zero, we created a null distribution for the dominance effect by calculating it on a randomized network where we shuffle the edges across pair of individuals (5000 times). We tested whether the observed effect is higher than the effect associated with the highest 0.05 of this null distribution. To investigate whether dominance is associated with higher influence strength, we performed dyadic regression (Tranmer et al. 2014; Hart et al. 2023) linear mixed model with peak cross-correlations as response, and individual dominance as predictor. We controlled for individual identity as a random effect. As fixed effects we included the sex of the individual and sex composition of the pair (male-male, male-female, female-female). To control for data dependencies by including individual identity in all our dyadic regressions, we use STAN and rstan (Stan Development Team, 2024). We ensured MCMC convergence by visually inspecting the chain and ensuring r-hat values were close to one.

### The consequences of social structure on rest

To test if closely ranked baboons are closer in space, we performed dyadic regression (Tranmer et al. 2014; Hart et al. 2023) by fitting a linear mixed model with pairwise median inter-individual distance as the response variable (standardized), individual dominance as the explanatory variable (standardized), individual identity as a random effect, and pair sex composition (male-male, male-female, female-female) as a control covariate. We investigated whether individuals that spend more time closer during the day influence each other’s behaviour more strongly during the night by performing dyadic regression where the log cross-correlation peak Ψ/Ψ_*max*_ (standardized) was the response (see methods 3.5 “Pairwise social influence”), log daytime spatial proximity (standardized) was the explanatory variable, individual identity as a random effect, and individual and pair sex composition (male-male, male-female, female-female) as a control covariate. Lastly, we investigated how dominance may impact network centrality and thus social rest disruption. To do this, we fitted two linear mixed models with centrality in the influence network and in the spatial proximity network, respectively, as response variables, individual dominance, and sex as predictor, and individual identity as random factor. We used eigenvector centrality as our network centrality metric, which assigns a high score to individuals connected to individuals that are also well connected to others (Csárdi et al. 2024). Eigen vector centrality *c*_*i*_ for individual *i* is the *i*^*th*^ entry of the eigenvector associated with the leading eigenvalue of the adjacency matrix of the network.

### Deposited data

Code available at: https://github.com/MarcoFele98/project_2_sleep

