## supplemental for "Dominant baboons experience more interrupted and less rest at night"

*[Figure S1: Baboons rest less during shorter nights.](#_Toc184897886)*	[2](#_Toc184897886)

*[Figure S3: Network of pairwise synchronization of rest](#_Toc184897888)*	[3](#_Toc184897888)

*[Figure S5: Probability of rest observed across the study](#_Toc184897894)*	[5](#_Toc184897894)

*[Figure S6: Defining rest state.](#_Toc184897895)*	[8](#_Toc184897895)

### SUPPLEMENTARY FIGURES - RESULTS

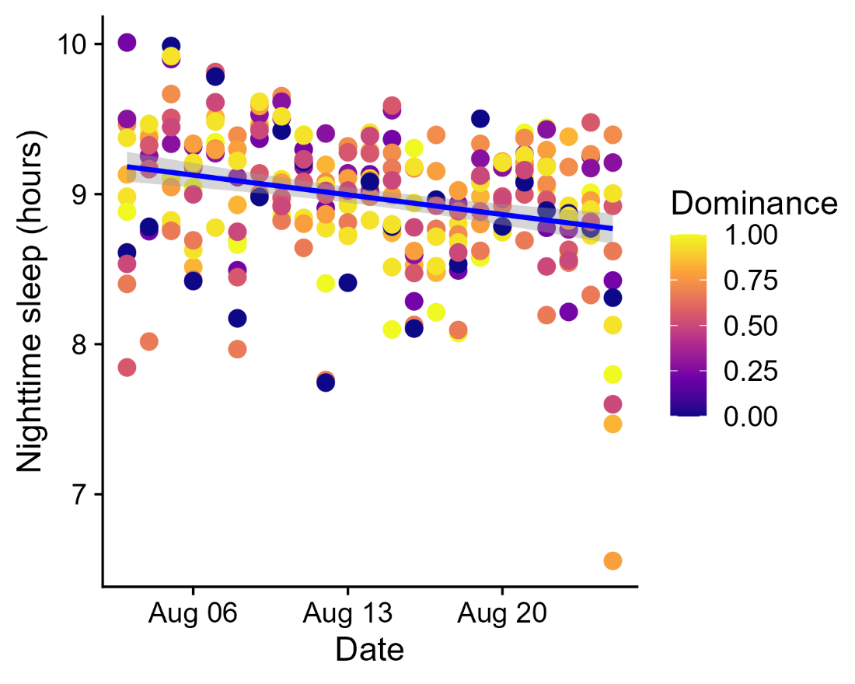

Figure S1: Baboons rest less during shorter nights. Nighttime rest (hours) over the study period moving from Winter (August) to Spring (September). Each datapoint represents the number hours one individual spent resting for one night, and the colour of the point is the individual’s dominance rank scaled between 0 (least dominant) and 1 (most dominant).

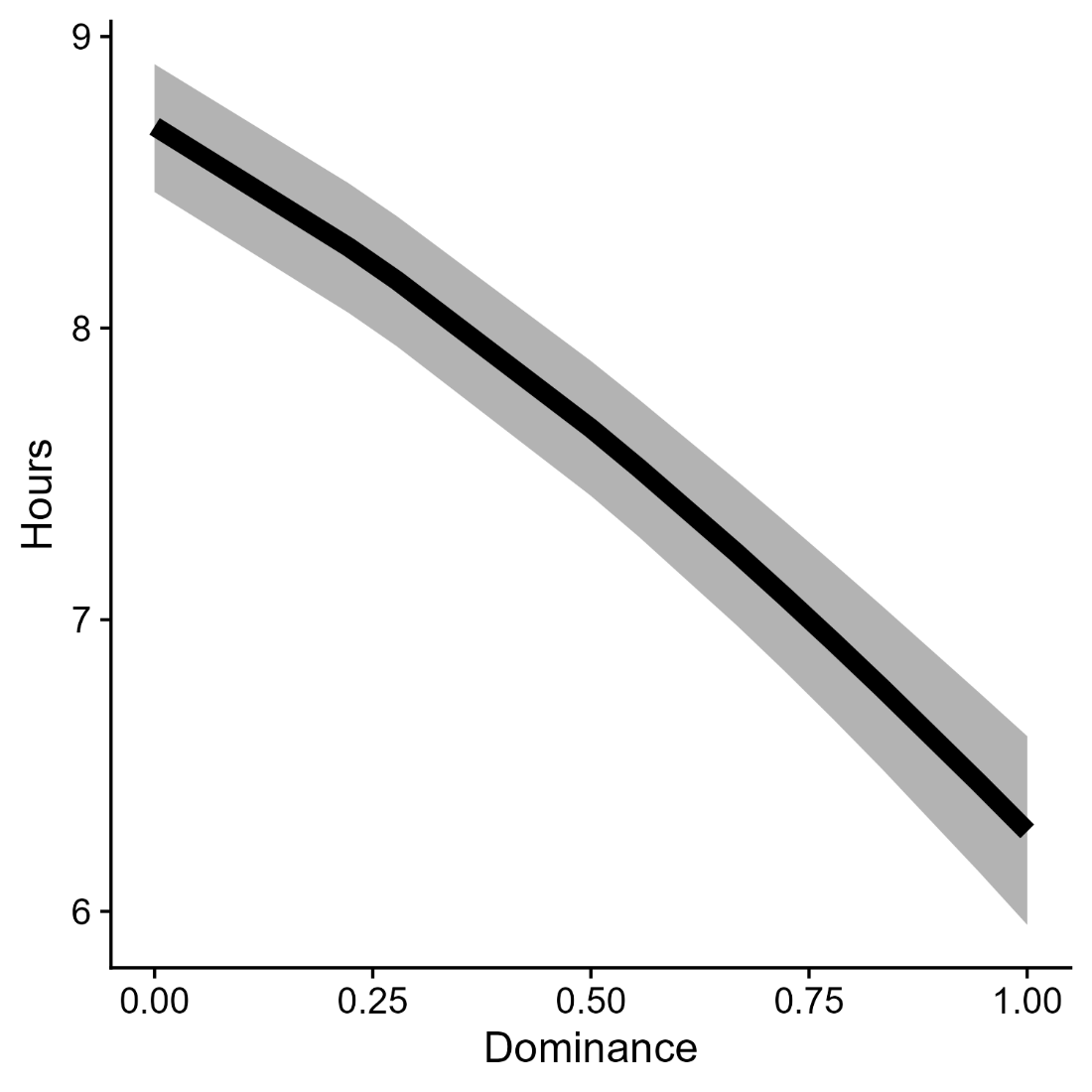

Figure S2: Dominant baboons rest less. Markov chain model predictions for the time baboons spend resting as a function of dominance. Predicted effect is shown for an individual with dominance 0.5, and when no other individual is resting.

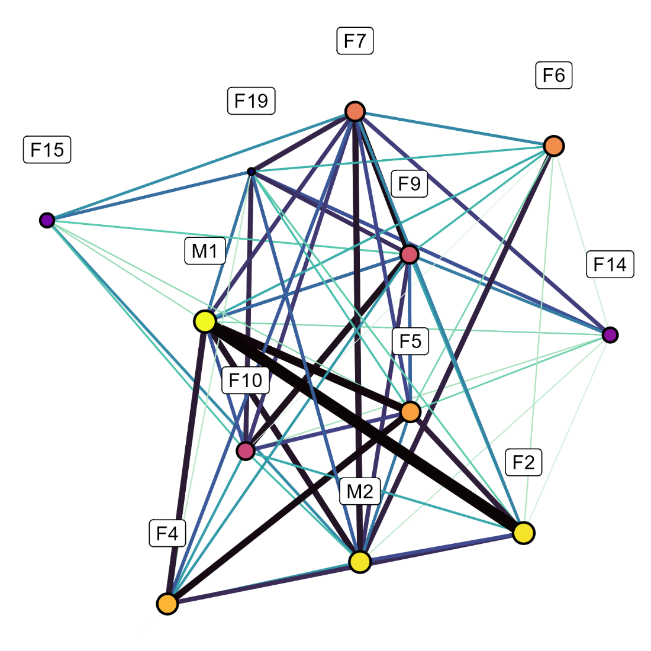

Figure S3: Network of pairwise synchronization of rest**.** Each baboon is a node (brighter colour and bigger size indicating higher dominance), and the edge width (and darker colours) indicates greater synchrony between a pair. Synchrony (edges) represents how likely it is that the two individuals are in the same state compared to if they behaved independently. Data available for n=12 individuals simultaneously from acceleration data (see Methods for full details).

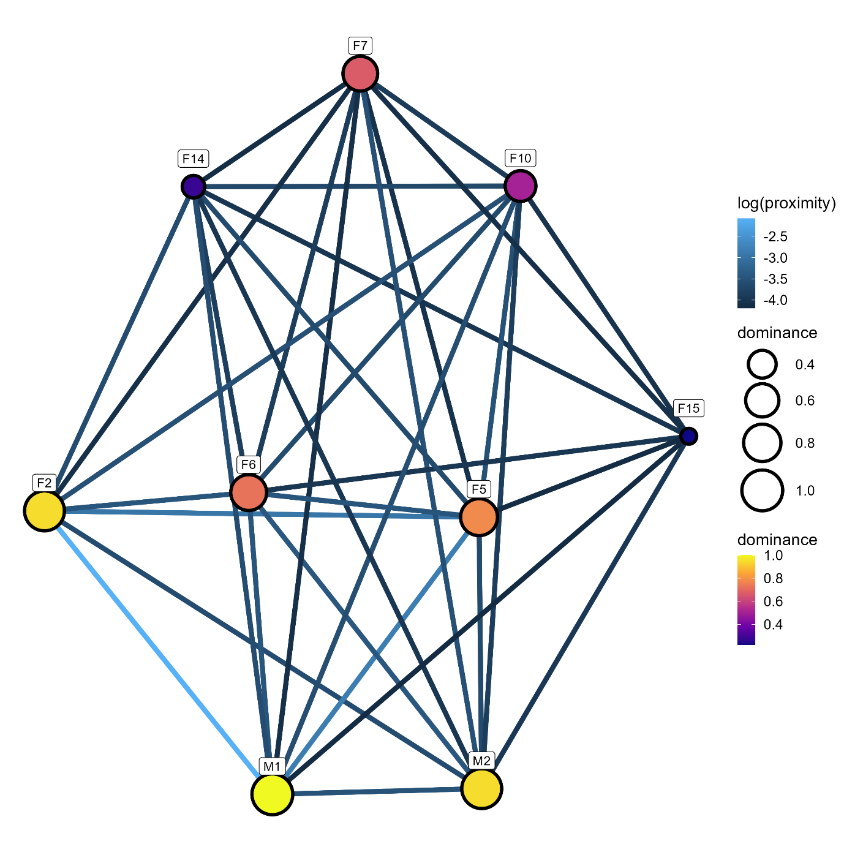

Figure S4. Spatial network based on individual median pairwise distance**.** Each baboon is a node (brighter colour and bigger size indicating higher dominance), and the edge colour indicates closer proximity (median distance) between a pair (lighter colour closer). Data available from simultaneous GPS for n=9 individuals (see Methods for full details).

### SUPPLEMENTARY TABLES - RESULTS

Table S1: Predictors of wakefulness. Predicted effects of dominance, number of others resting (and their interaction), on wakefulness, controlling controlled for the effects the previous night’s rest, night-time rainfall, temperature, and moon illumination. Random effects are individual and night ID, to control for non-independence of data for individuals within nights.

| coefficent | estimate | standard error | p.value |
| --- | --- | --- | --- |
| intercept | -3.85 | 2.56 | 0.13 |
| dominance | -1.85 | 0.1 | 0 |
| avegare dominance of resting individuals | -1.31 | 0.07 | 0 |
| number rest exlcuding focal individual | 0.09 | 0 | 0 |
| night duration | 0 | 0 | 0.4 |
| rest in previous night | 0.27 | 0 | 0 |
| did it rain | 0.35 | 0.07 | 0 |
| average temperature | -0.04 | 0.02 | 0 |
| fraction of moon visible | 0.19 | 0.12 | 0.01 |
| interaction domince with average dominance of resting individuals | 2.5 | 0.12 | 0 |
| interaction dominance with number of resting individuals | 0.04 | 0 | 0 |
| random effect id | 0.07 |  |  |
| random effect night | 0.14 |  |  |

Table S2: Predictors of rest quality. Predicted effects of dominance, number of others resting (and their interaction), on rest quality, controlling controlled for the effects the previous night’s rest, night-time rainfall, temperature, and moon illumination. Random effects are individual and night ID, to control for non-independence of data for individuals within nights.

| coefficent | estimate | standard error | p.value |
| --- | --- | --- | --- |
| intercept | -4.29 | 5.96 | 0.47 |
| dominance | 1.65 | 0.18 | 0 |
| avegare dominance of resting individuals | 1.22 | 0.08 | 0 |
| number rest exlcuding focal individual | -0.16 | 0 | 0 |
| night duration | 0 | 0 | 0.87 |
| rest in previous night | -0.55 | 0 | 0 |
| did it rain | -0.29 | 0.167 | 0.08 |
| average temperature | 0.06 | 0.04 | 0.12 |
| fraction of moon visible | -0.48 | 0.3 | 0.1 |
| interaction domince with average dominance of resting individuals | -2.41 | 0.11 | 0 |
| interaction dominance with number of resting individuals | 0 | 0 | 0.89 |
| random effect id | 0.17 |  |  |
| random effect night | 0.33 |  |  |

### SUPPLEMENTARY FIGURES - METHODS

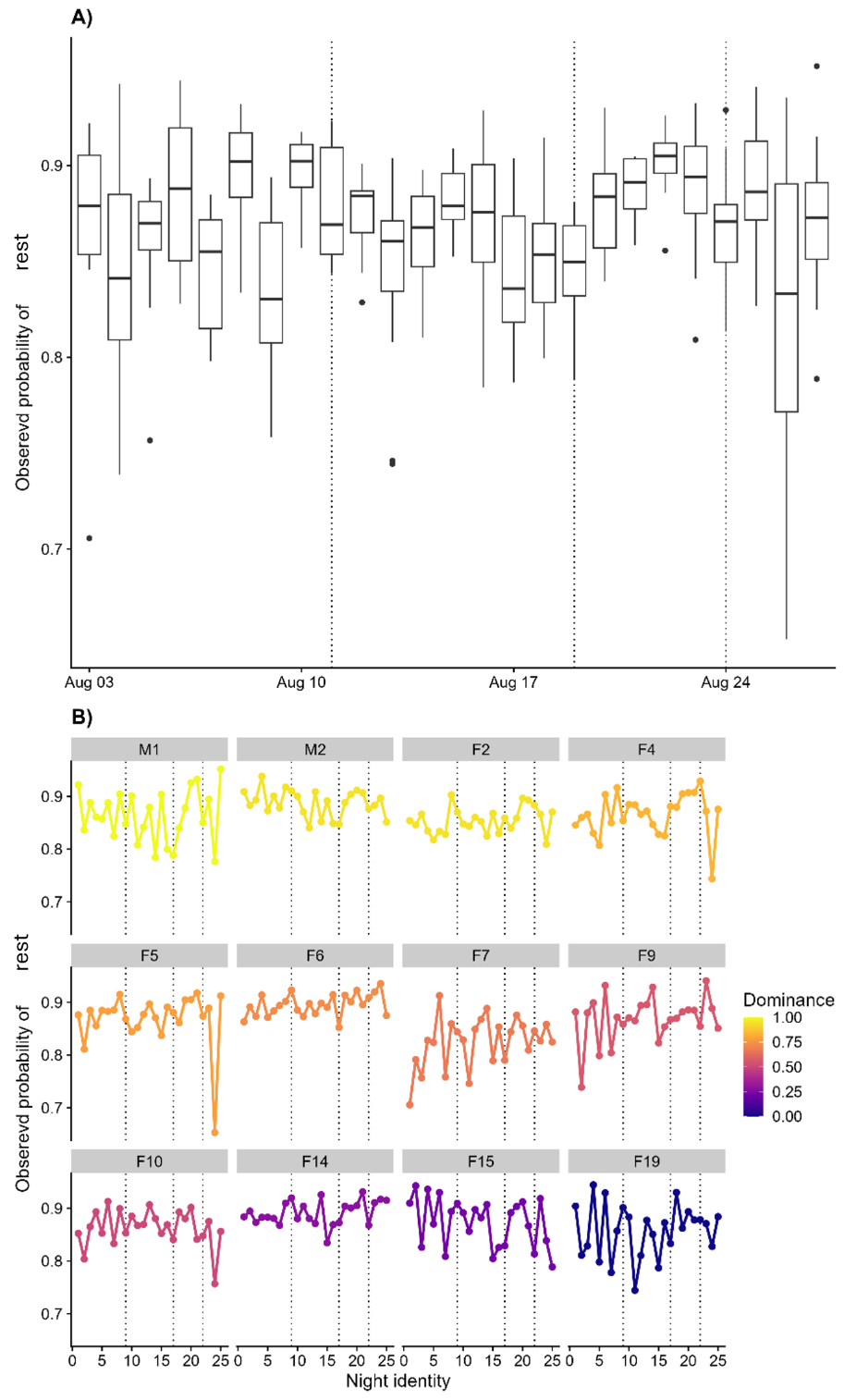

Figure S5: Probability of rest observed across the study**.** A) Boxplot of observed rest probability across all individuals within each study night. B) Time series of the observed rest probability for every individual. In both A and B, the vertical lines correspond to nights in which the troop slept in a natural space, and other nights the baboons slept in urban space.

(the following figure S6 has 12 panels, one for baboon; figure legend at bottom of figure)

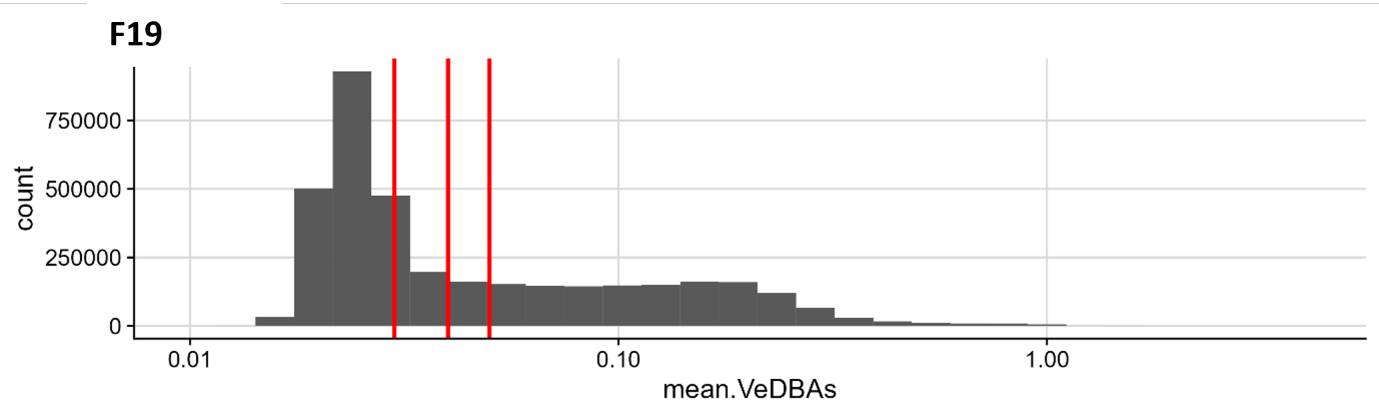

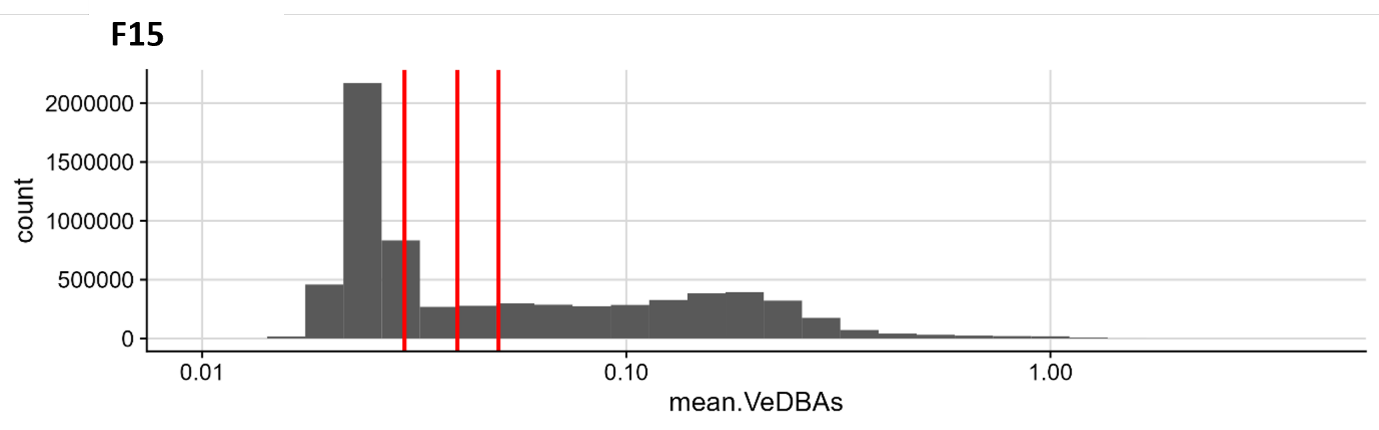

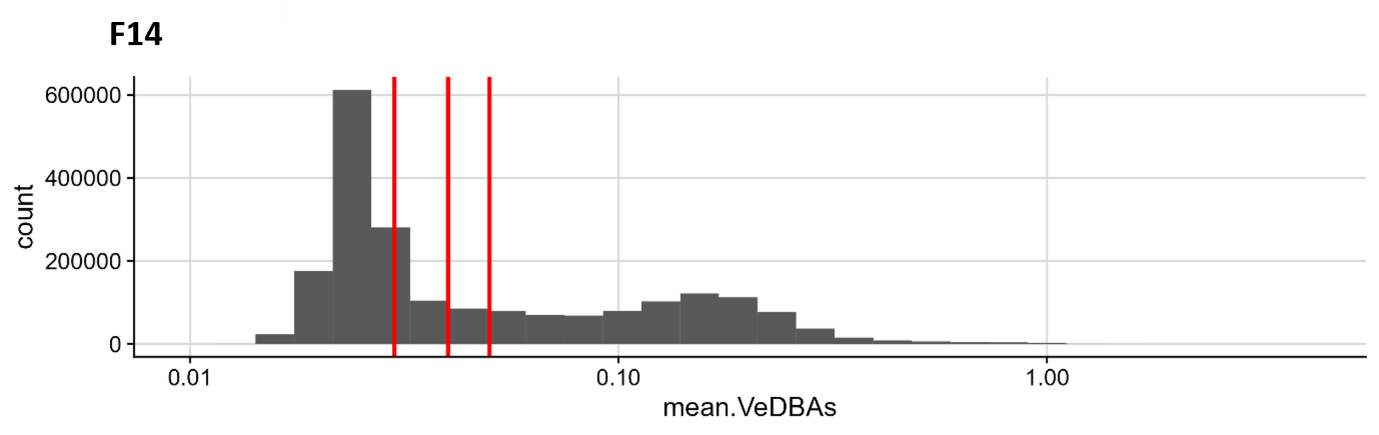

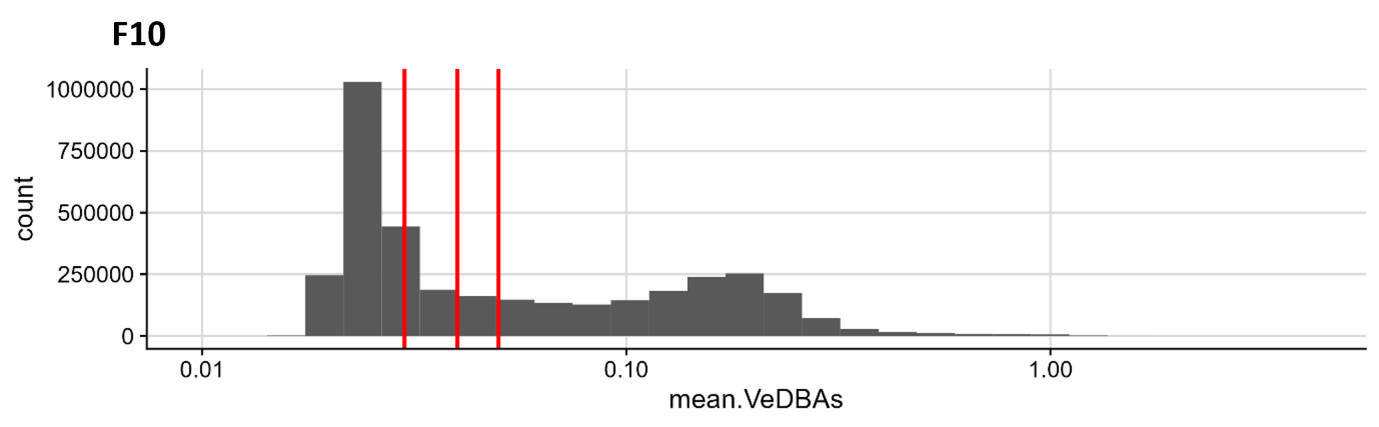

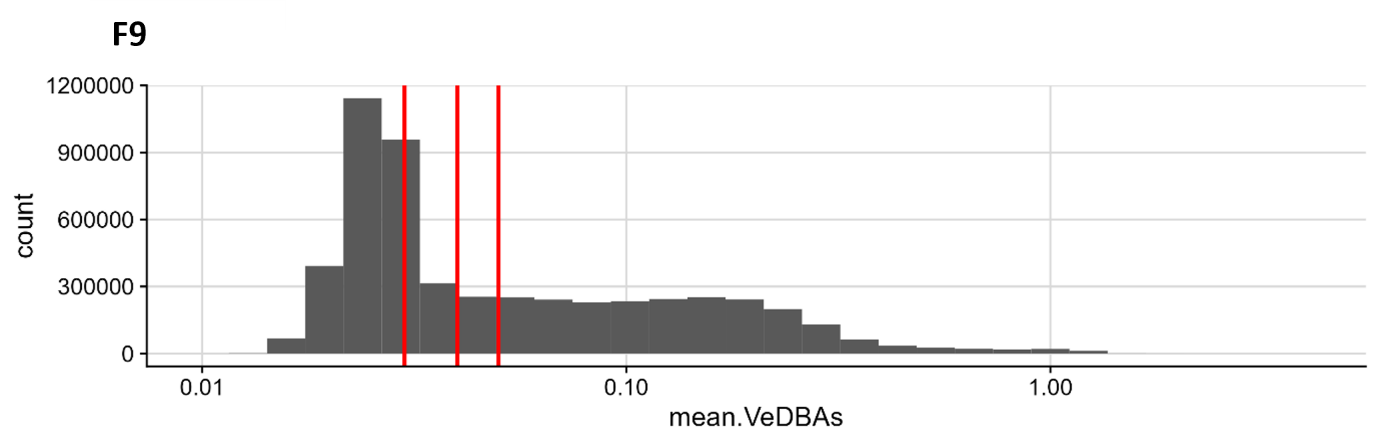

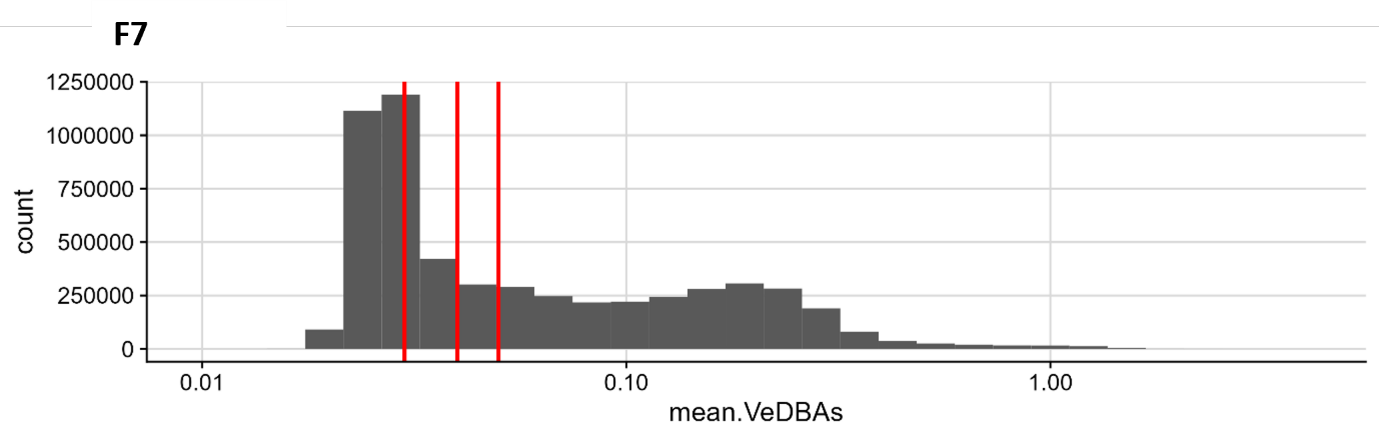

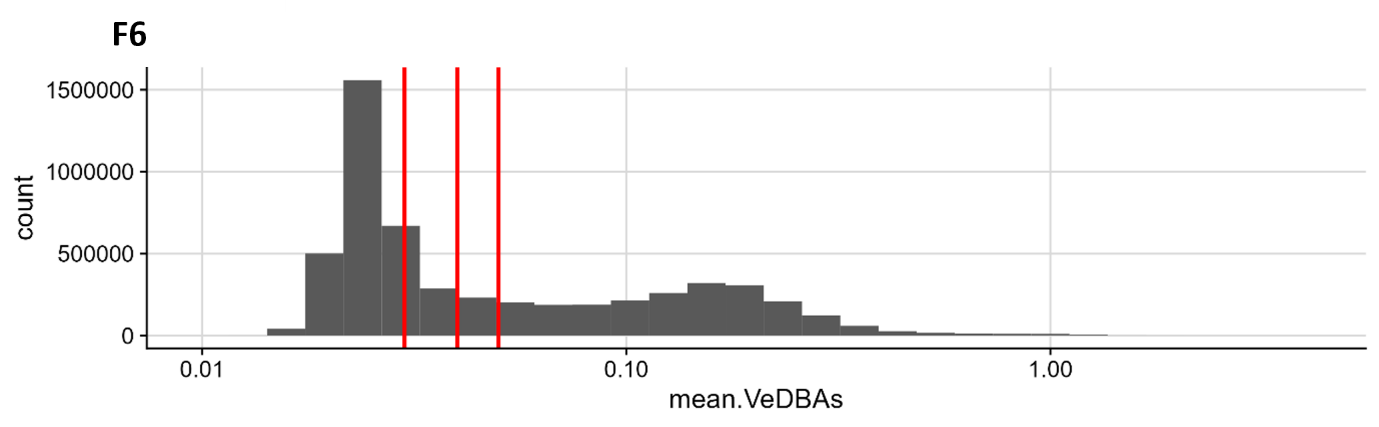

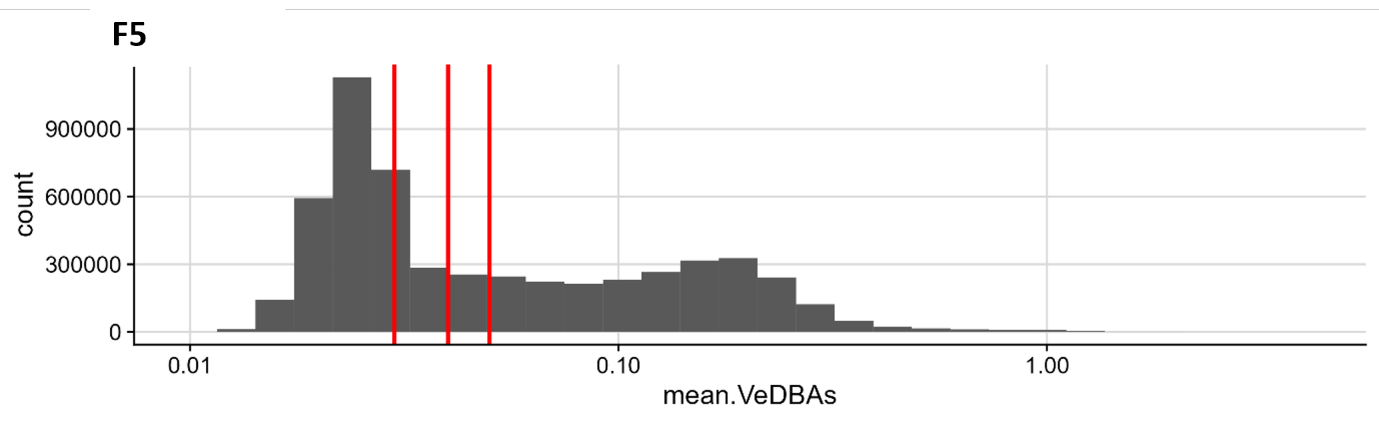

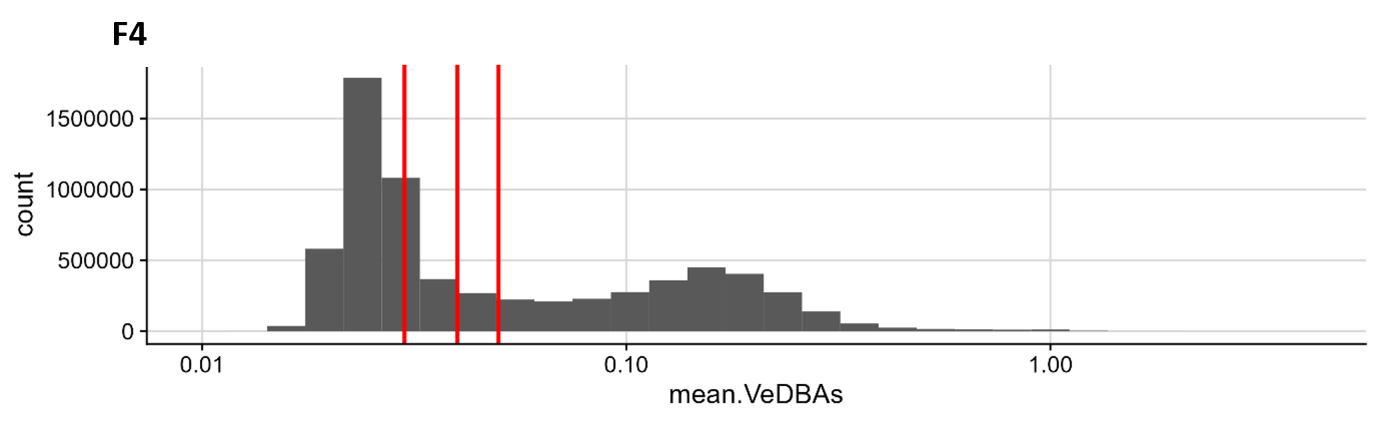

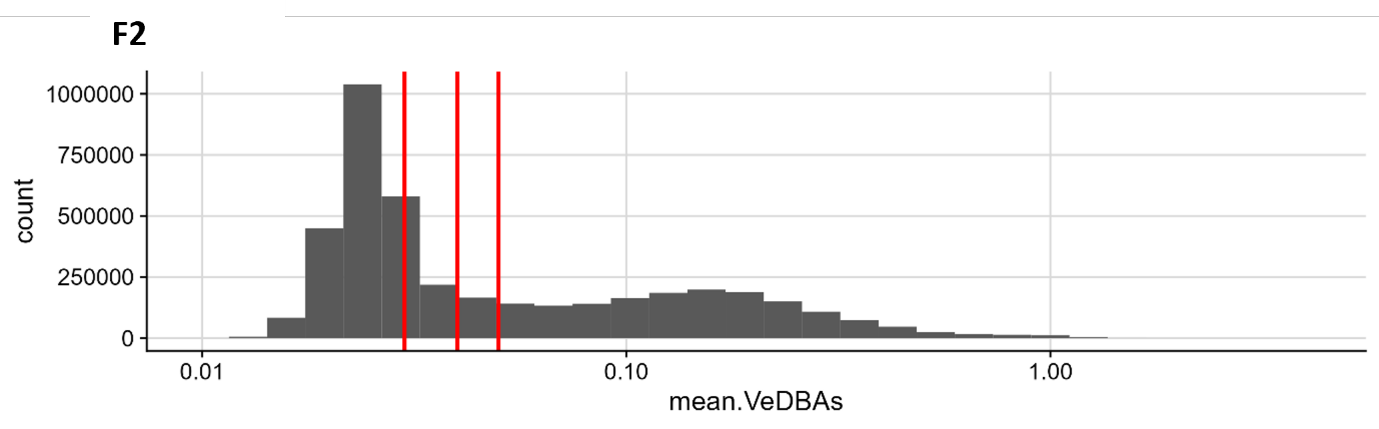

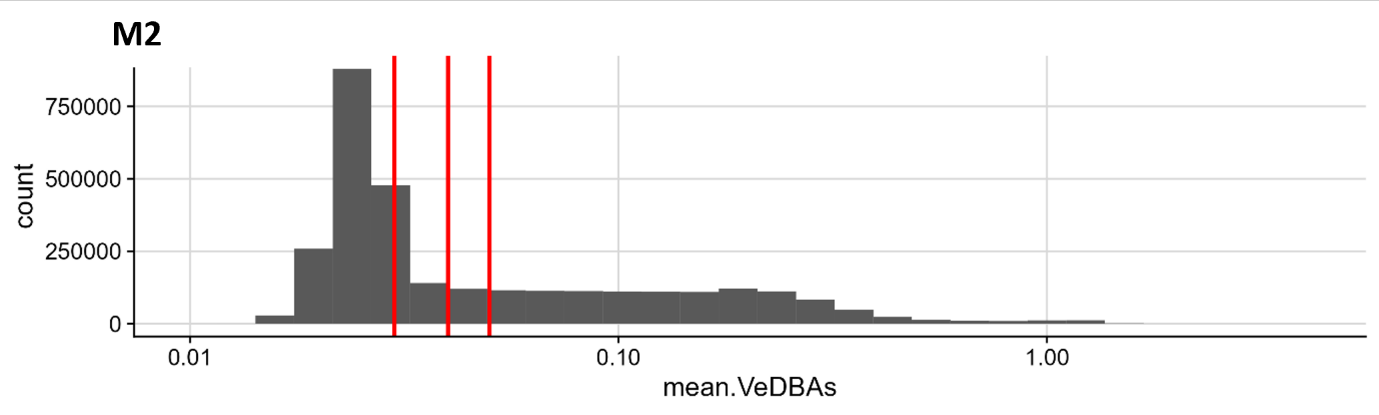

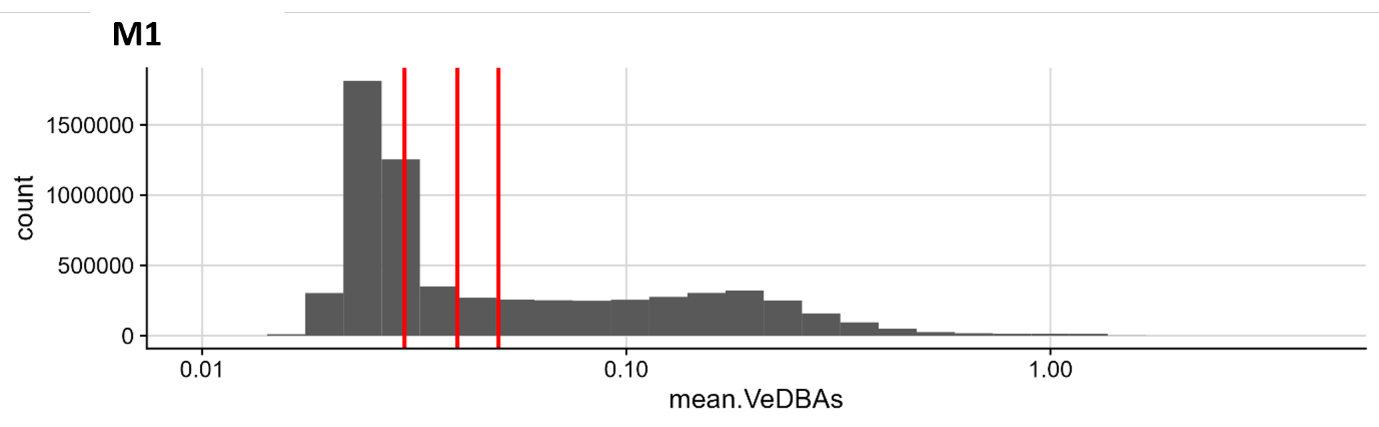

Figure S6: Defining rest state. Distribution of acceleration values (VeDBA) for every individual. x axis is in log scale. Vertical lines show the different thresholding values (0.3, 0.4, 0.5) used to identify rest and infer rest state for our sensitivity analyses (Tables S6-S9).

### SUPPLEMENTARY TABLES - METHODS

Table S3: Collar GPS data. GPS data periods per individual are given. We have some daytime GPS data between 30/7/2018 and 15/9/2018 for 13 individuals.

| ID | start | end |
| --- | --- | --- |
| F15 | 7/30/2018 | 9/7/2018 17:31 |
| F18 | 7/30/2018 11:15 | 9/7/2018 17:31 |
| F13 | 8/2/2018 12:30 | 9/7/2018 17:31 |
| F2 | 7/30/2018 11:15 | 9/7/2018 17:31 |
| M1 | 7/30/2018 11:15 | 9/7/2018 17:31 |
| F6 | 7/30/2018 11:15 | 9/7/2018 9:43 |
| F14 | 8/2/2018 12:30 | 8/25/2018 17:59 |
| F5 | 7/30/2018 11:15 | 9/7/2018 17:31 |
| F9 | 7/30/2018 11:15 | 8/15/2018 10:48 |
| F7 | 7/30/2018 11:15 | 9/7/2018 17:31 |
| M2 | 7/30/2018 11:15 | 9/7/2018 17:31 |
| F17 | 8/2/2018 11:00 | 9/7/2018 17:31 |
| F10 | 7/30/2018 11:15 | 9/7/2018 17:31 |

Table S4: Collar accelerometer data**.** Acceleration data periods per individual are given. We have full nighttime data between 2nd August 2018 and 27th of August 2018 for 12 individuals.

| baboon | start | date |
| --- | --- | --- |
| F15 | 7/27/2018 | 10/15/2018 23:59 |
| F4 | 7/26/2018 | 10/12/2018 23:59 |
| F2 | 7/26/2018 | 9/11/2018 23:59 |
| M1 | 7/31/2018 | 10/14/2018 23:59 |
| F6 | 7/27/2018 | 9/27/2018 23:59 |
| F14 | 8/3/2018 | 8/26/2018 23:59 |
| F5 | 7/27/2018 | 9/27/2018 23:59 |
| F9 | 7/27/2018 | 9/26/2018 23:59 |
| F7 | 7/27/2018 | 9/29/2018 23:59 |
| F19 | 7/31/2018 | 9/10/2018 23:59 |
| M2 | 7/31/2018 | 9/3/2018 23:59 |
| F10 | 7/31/2018 | 9/11/2018 1:58 |

Table S5. Astronomical night. Defined as when the geometric centre of the sun is 18 degrees below the horizon, occurring approximately between ~20:00 and ~06:00 in our study system. We obtained this data with the R package “suncalc” (Thieurmel and Elmarhraoui 2017).

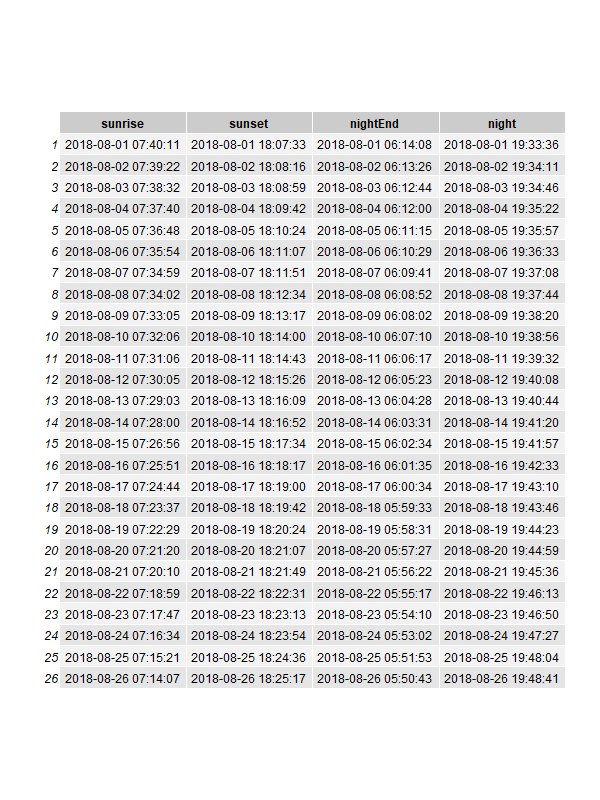

Table S6: Markov chain model results for wakefulness. 0.03 VeDBA as threshold.

| coefficent | estimate | standard error | p.value |
| --- | --- | --- | --- |
| intercept | -3.85 | 2.55 | 0.13 |
| dominance | -1.85 | 0.1 | 0 |
| avegare dominance of resting individuals | -1.31 | 0.07 | 0 |
| number rest exlcuding focal individual | 0.09 | 0 | 0 |
| night duration | 0 | 0 | 0.43 |
| rest in previous night | 0.27 | 0 | 0 |
| did it rain | 0.35 | 0.07 | 0 |
| average temperature | -0.5 | 0.02 | 0.01 |
| fraction of moon visible | 0.19 | 0.12 | 0.13 |
| interaction domince with average dominance of resting individuals | 2.5 | 0.1 | 0 |
| interaction dominance with number of resting individuals | 0.04 | 0 | 0 |
| random effect id | 0.07 |  |  |
| random effect night | 0.14 |  |  |

Table S7: Markov chain model results for res quality. 0.03 VeDBA as threshold.

| coefficent | estimate | standard error | p.value |
| --- | --- | --- | --- |
| intercept | -4.28 | 5.96 | 0.47 |
| dominance | 1.65 | 0.18 | 0 |
| avegare dominance of resting individuals | 1.22 | 0.08 | 0 |
| number rest exlcuding focal individual | -0.16 | 0 | 0 |
| night duration | 0 | 0 | 0.87 |
| rest in previous night | -0.55 | 0 | 0 |
| did it rain | -0.29 | 0.17 | 0.08 |
| average temperature | 0.06 | 0.04 | 0.12 |
| fraction of moon visible | -0.48 | 0.3 | 0.1 |
| interaction domince with average dominance of resting individuals | -2.41 | 0.11 | 0 |
| interaction dominance with number of resting individuals | 0 | 0 | 0.9 |
| random effect id | 0.17 |  |  |
| random effect night | 0.33 |  |  |

Table S8: Markov chain model results for wakefulness. 0.05 VeDBA as threshold.

| coefficent | estimate | standard error | p.value |
| --- | --- | --- | --- |
| intercept | -4.7 | 6.14 | 0.13 |
| dominance | -2.39 | 0.17 | 0 |
| avegare dominance of resting individuals | -1.4 | 0.14 | 0 |
| number rest exlcuding focal individual | 0.4 | 0 | 0 |
| night duration | 0 | 0 | 0.43 |
| rest in previous night | 0.41 | 0.01 | 0 |
| did it rain | 0.3 | 0.13 | 0 |
| average temperature | -0.05 | 0.03 | 0.01 |
| fraction of moon visible | 0.3 | 0.26 | 0.13 |
| interaction domince with average dominance of resting individuals | 2.1 | 0.21 | 0 |
| interaction dominance with number of resting individuals | 0.6 | 0 | 0 |
| random effect id | 0.1 |  |  |
| random effect night | 0.24 |  |  |

Table S9: Markov chain model results for rest quality. 0.05 VeDBA as threshold.

| coefficent | estimate | standard error | p.value |
| --- | --- | --- | --- |
| intercept | -9.37 | 12.3 | 0.45 |
| dominance | 4.13 | 0.17 | 0 |
| avegare dominance of resting individuals | 2.83 | 0.14 | 0 |
| number rest exlcuding focal individual | -0.23 | 0 | 0 |
| night duration | 0 | 0 | 0.68 |
| rest in previous night | -0.99 | 0.13 | 0 |
| did it rain | -0.48 | 0.34 | 0.16 |
| average temperature | 0.11 | 0.08 | 0.16 |
| fraction of moon visible | -0.82 | 0.6 | 0.17 |
| interaction domince with average dominance of resting individuals | -5.59 | 0.2 | 0 |
| interaction dominance with number of resting individuals | -0.07 | 0 | 0 |
| random effect id | 0.12 |  |  |
| random effect night | 0.69 |  |  |

### SUPPLEMENTARY METHODS

#### Full model specification

$${state}_{t+1}\sim bernoulli\left( p_{S\to S} | {state}_{t}=1 \right)$$

${state}_{t+1}\sim bernoulli\left( 1-p_{A\to A} | {state}_{t}=0 \right)$

$$\mathrm{with}p\text{:}\text{ }p_{S\to S},p_{A\to A}$$

$$logit\left( p \right)\text{ }\text{=}\text{ }\text{(1|id)}\text{+}\text{sleep\_previous\_night}\text{+}\text{dominance}\text{*(}\text{number\_sleeping}\text{+}$$

$$\text{avg\_dominance\_sleeping}\text{)+}$$

$$\text{night\_duration+(1|night)+mean\_temperature}\text{+}\text{precipitation}\text{+}\text{moon\_fraction}$$

Complete specification of Markov chain model. Each parameter is estimated through a logistic regression, considering individual characteristics (green), social environment (red), and control covariates (purple). State = 1 corresponds to resting, state = 0 corresponds to active.

#### Calculating rest probability and its confidence interval from transition parameters

Given the Markov chain model described in the main text, we have a transition matrix ***M*** which for convenience we parametrize by considering the transition probabilities between states $p_{S\to A}$and $p_{A\to S}$as the free parameters estimated by the logistic regression models.

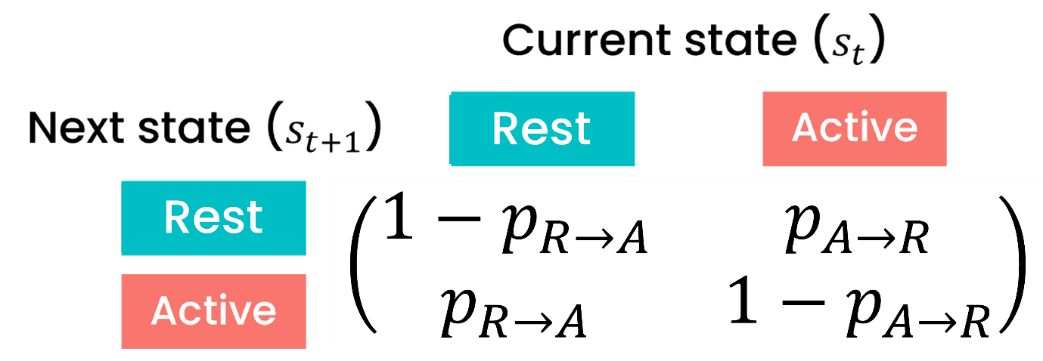

Figure 2: Transition matrix for the Markov chain model.

To find the rest probability $p_{S}$, we consider $\boldsymbol{v}_{\boldsymbol{t}}$ the column vector of size 2 which defines the probability of rest (first entry) or active (second entry) at time t. We can find $\boldsymbol{v}_{\boldsymbol{t}}$by repeatedly multiplying the transition matrix ***M*** (fig) with the initial probabilities $\boldsymbol{v}_{\boldsymbol{0}}$ so that $\boldsymbol{v}_{\boldsymbol{t}}$ =$\boldsymbol{M}^{\boldsymbol{t}} \boldsymbol{v}_{\boldsymbol{0}}$. We can find the steady state, i.e., the state probabilities that do not change between timesteps, by solving $\boldsymbol{v}_{\boldsymbol{t}\boldsymbol{+1}}$ = $\boldsymbol{M}\boldsymbol{v}_{\boldsymbol{t}}$. More explicitly we find the probability of rest $p_{S}$ that solves system of equations given by the dot product:

$$\binom{p_{S}}{p_{A}}=\left( \begin{matrix} 1-p_{S\to A} & p_{S\to A} \\ p_{A\to S} & 1-p_{S\to A} \end{matrix} \right).\binom{p_{S}}{p_{A}}$$

We find $p_{S}=\frac{p_{A\to S}}{p_{A\to S}+p_{S\to A}}$, which is then converted to the alternative parameterization in the main text. The intuition behind $p_{S}$ is that the probability of resting is equal to the “push” towards rest over the total “push”. The upper bound for the confidence interval of $p_{S}$is calculated by substituting $p_{A\to S}$ with $p_{A\to S}+SE(p_{A\to S})$ and $p_{S\to A}$ with $p_{S\to A}-SE(p_{S\to A})$, where SE(p) is the standard error of the parameter estimated from the logistic regression. The +/- signs are inverted to find the lower bound.

#### Validation of Markov chain model

In this section we address 1) the assumption that individual behaviour can be modelled as a Markov chain and 2) the assumption that the chain reached steady state, which we use to find the probability of resting $p_{S}$.

The Markov chain model does not capture every aspect of nighttime rest (Figure S7 & S8). For example, we cannot model the long tails of the frequency distribution of duration of bouts of continuous behaviour because our Markov chain model results in an exponential distribution for behavioural duration, underrepresenting long bouts of continuous behaviour, which are very rare but present in the data. This could be avoided by using a semi-Markov process, where the time spent in state is included as covariate predicting the state transition probabilities, but is not necessary to understand the patterns of rest we investigated here and is outside the scope of our project.

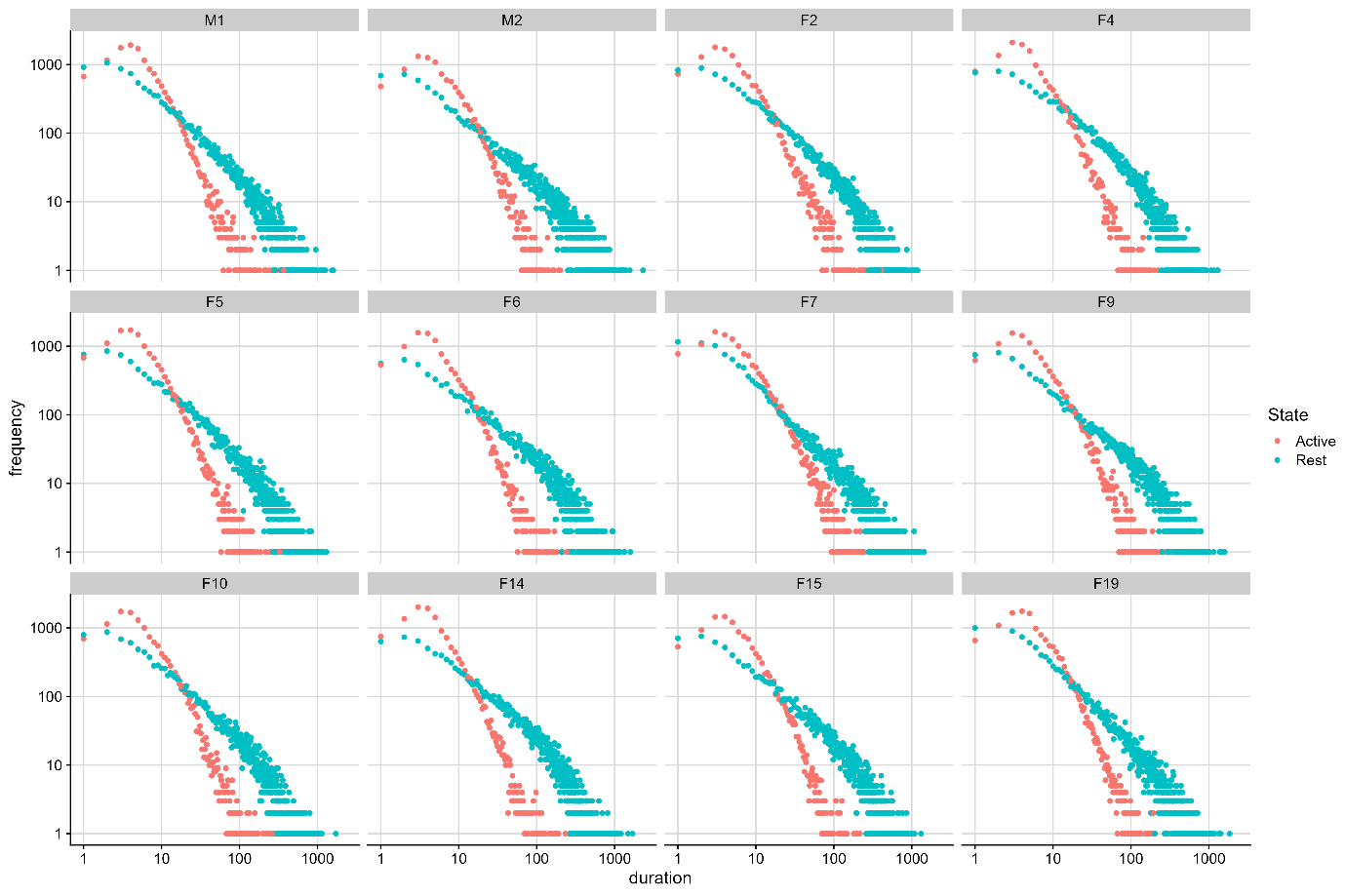

**Figure S7: Frequency distribution of duration of active and rest bouts from our raw data.** Frequency of active and rest in our raw data, for each baboon.

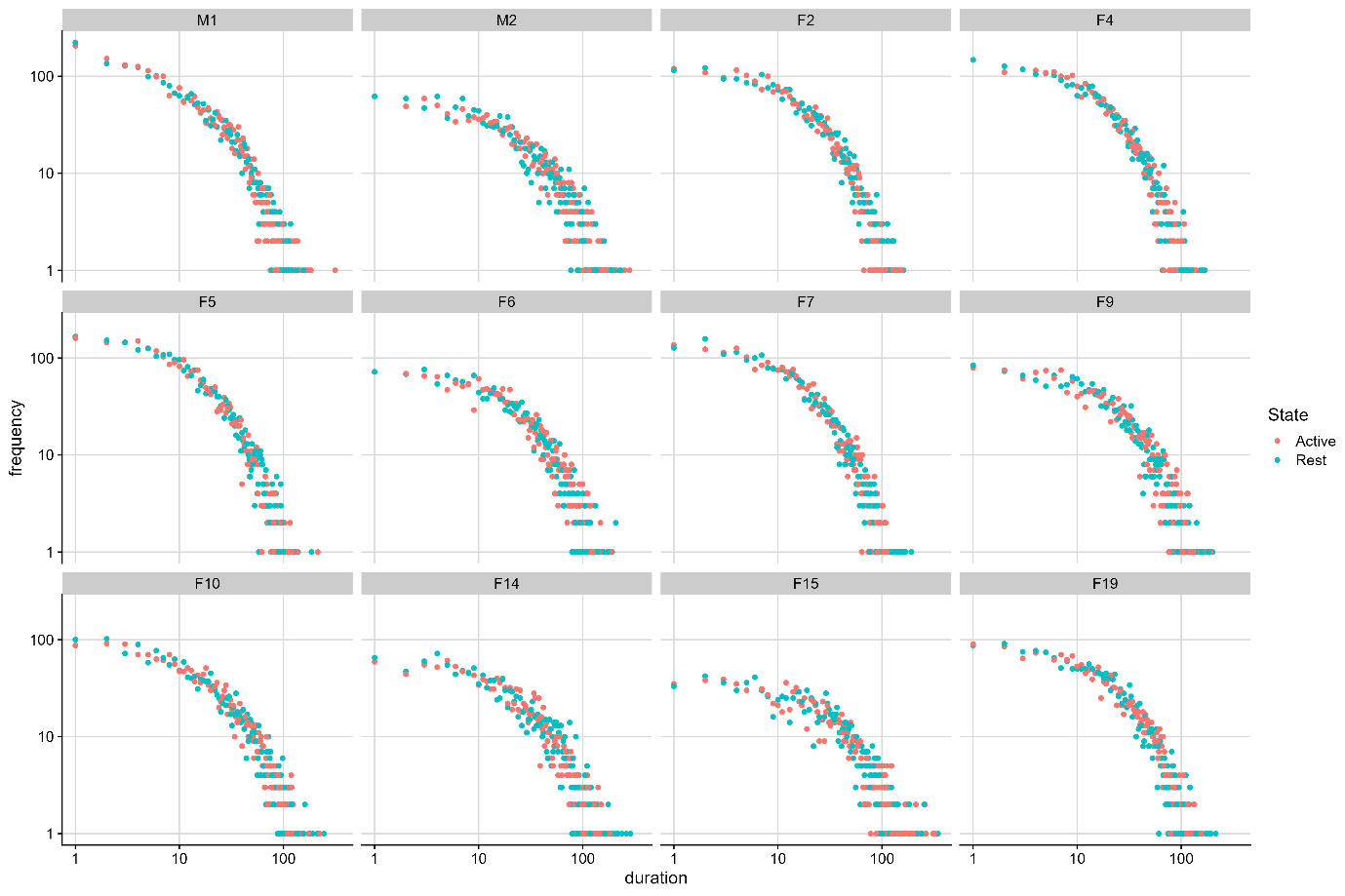

**Figure S8: Frequency distribution of duration of active and rest bouts from our model.** Frequency of active and rest periods as predicted by our Markov chain model, for each baboon.

Regarding the assumption that the chain reached steady state used to calculate $p_{S}$, we check whether assuming the chain reached steady state gives wrong predictions, regardless of whether the steady state assumption holds or not. We can do so by comparing the predicted time an individual spent resting according to our Markov chain model, which assumes steady state, with the observed time found in the data. We find model predictions by calculating the predicted proportion of time an individual spends resting for every possible combination of explanatory variables in the data. For every individual we then marginalize by averaging the predicted time spent resting over all possible combinations of covariates, accounting for different number of instances a specific combination of covariates occurred by weighting the probability of resting according to the number of observations. Regardless of whether the steady state assumption holds, our model predicts the proportion of time an individual spends resting within an error of ±0.003, which corresponds to a total maximum prediction error of ±0.6 hours out of 203 total rest hours across 25 days. We show the performance of our model in the same way marginalizing for different sets of explanatory variables (Figure S9-S15).

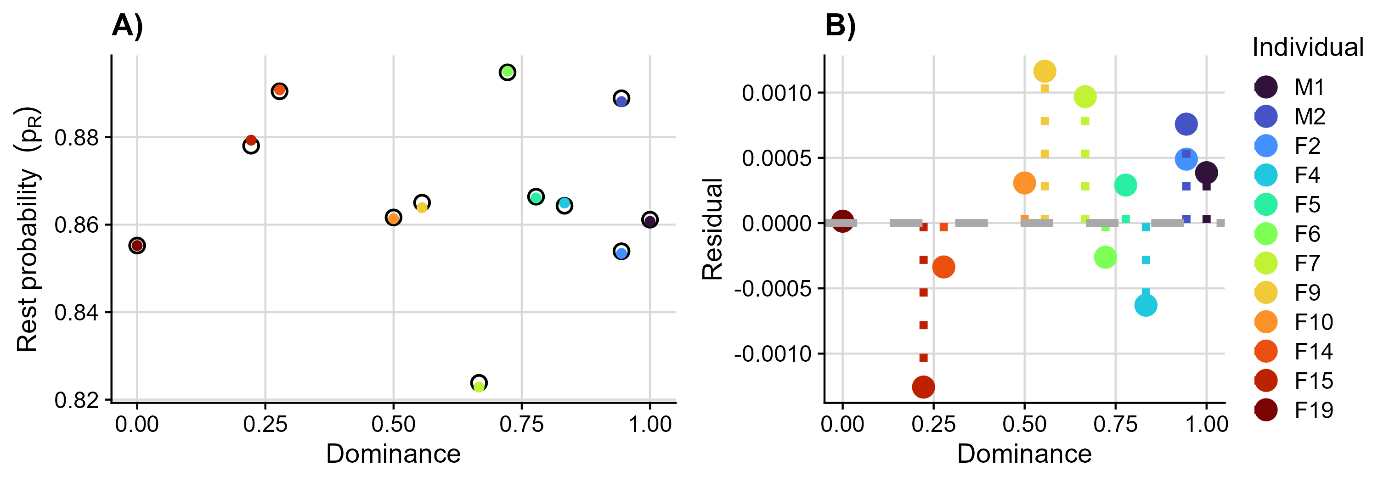

**Figure S9. Model performance.** Performance of our model for rest probability and residual of rest probability for study subjects.

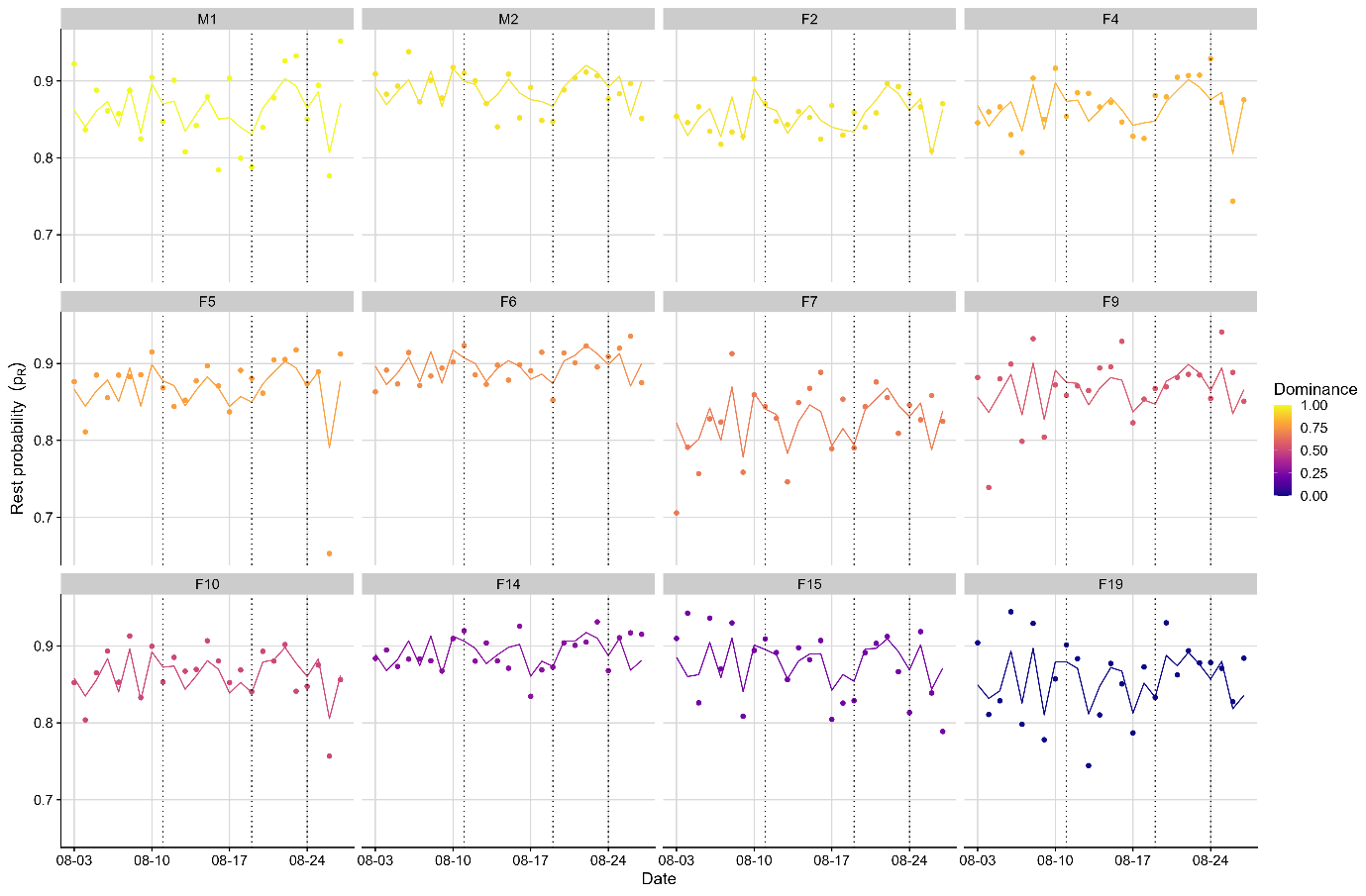

Figure S10. Model predicted rest probabilities (lines) compared to raw data (points) for each night. Every graph is an individual.

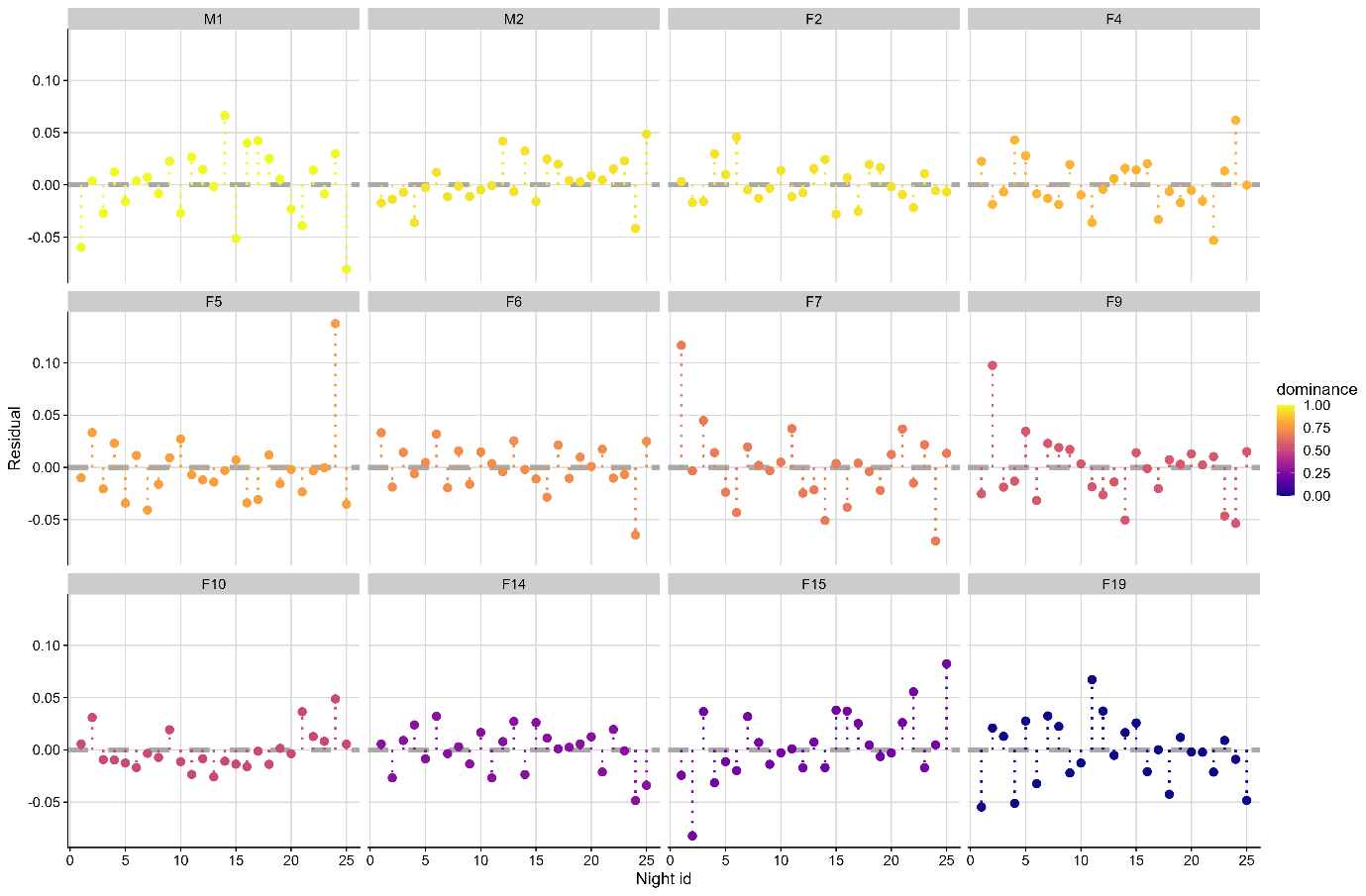

Figure S11. Model residuals (predictions minus observed value) for each night. Every graph is an individual.

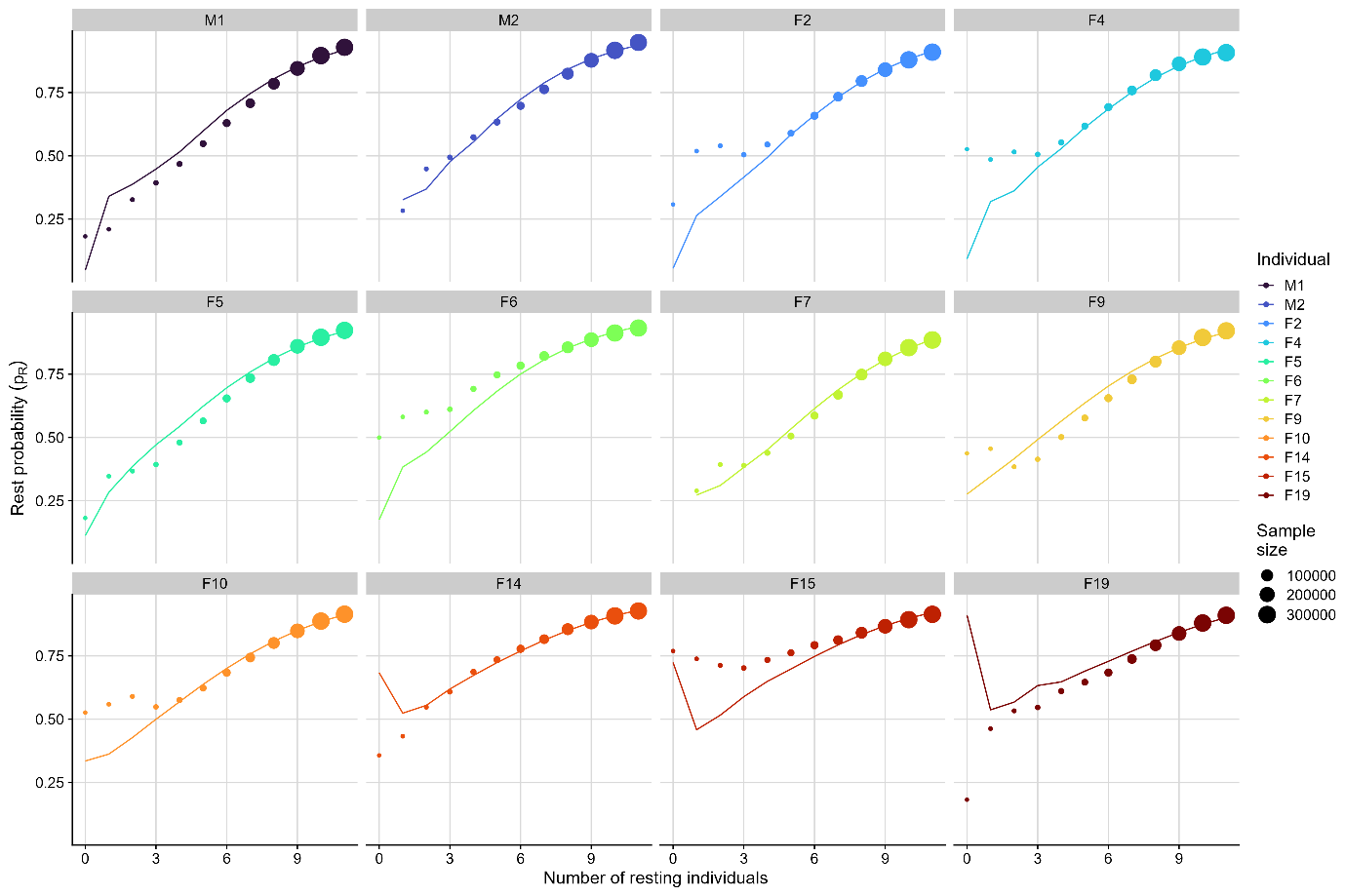

Figure S12. Model predicted rest probability (lines) compared to raw data (points, size is sample size) for the number of individuals simultaneously resting. Every graph is one individual.

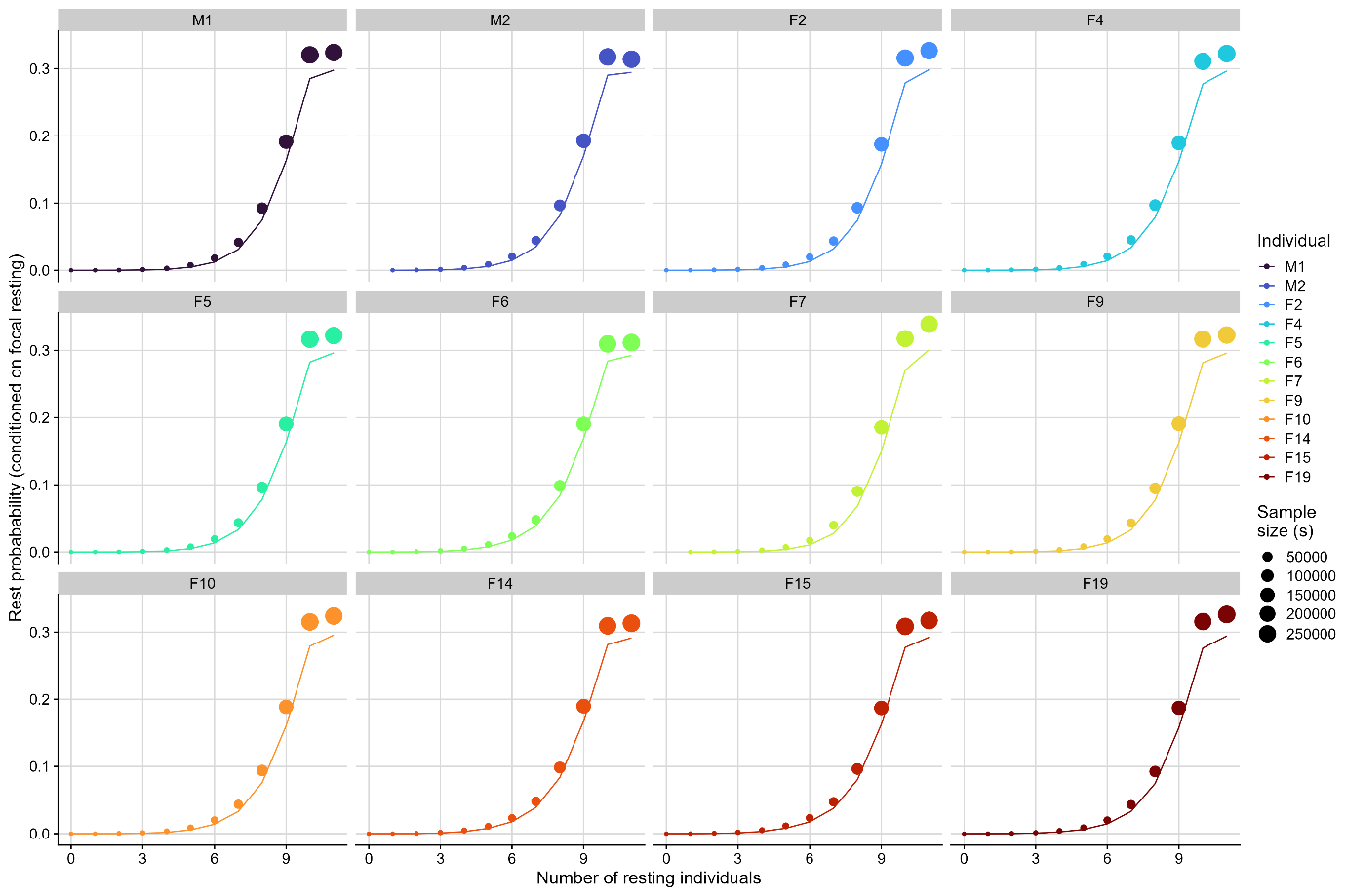

Figure S13. Model predicted rest probability (lines) compared to raw data (points, size is sample size) for the number of individuals simultaneously resting when conditioning on the focus individual also resting. Every graph is one individual.

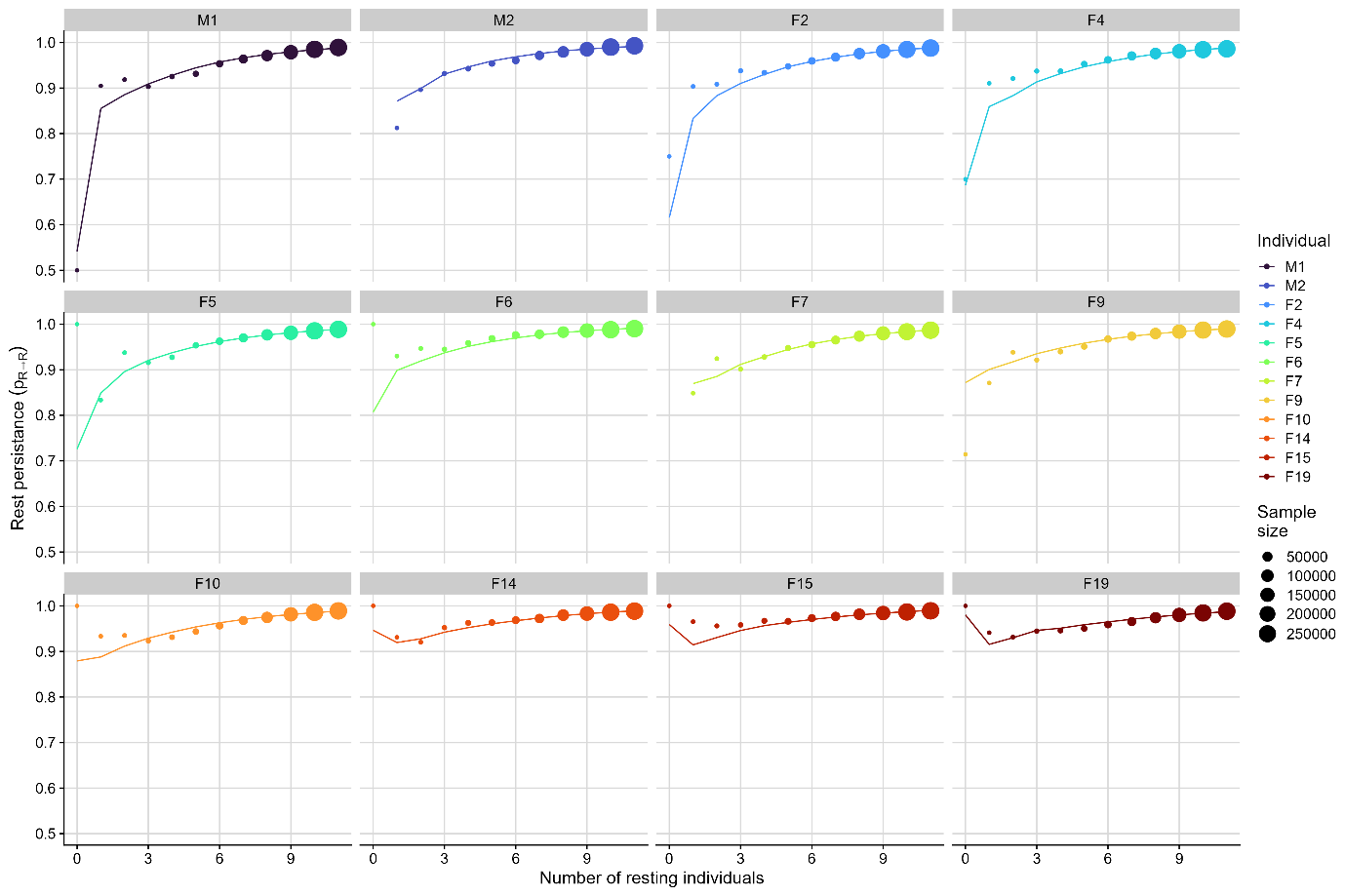

Figure S14. Model predicted rest quality (lines) compared to raw data (points, size is sample size) for the number of individuals simultaneously resting when conditioning on the focus individual also resting. Every graph is one individual.

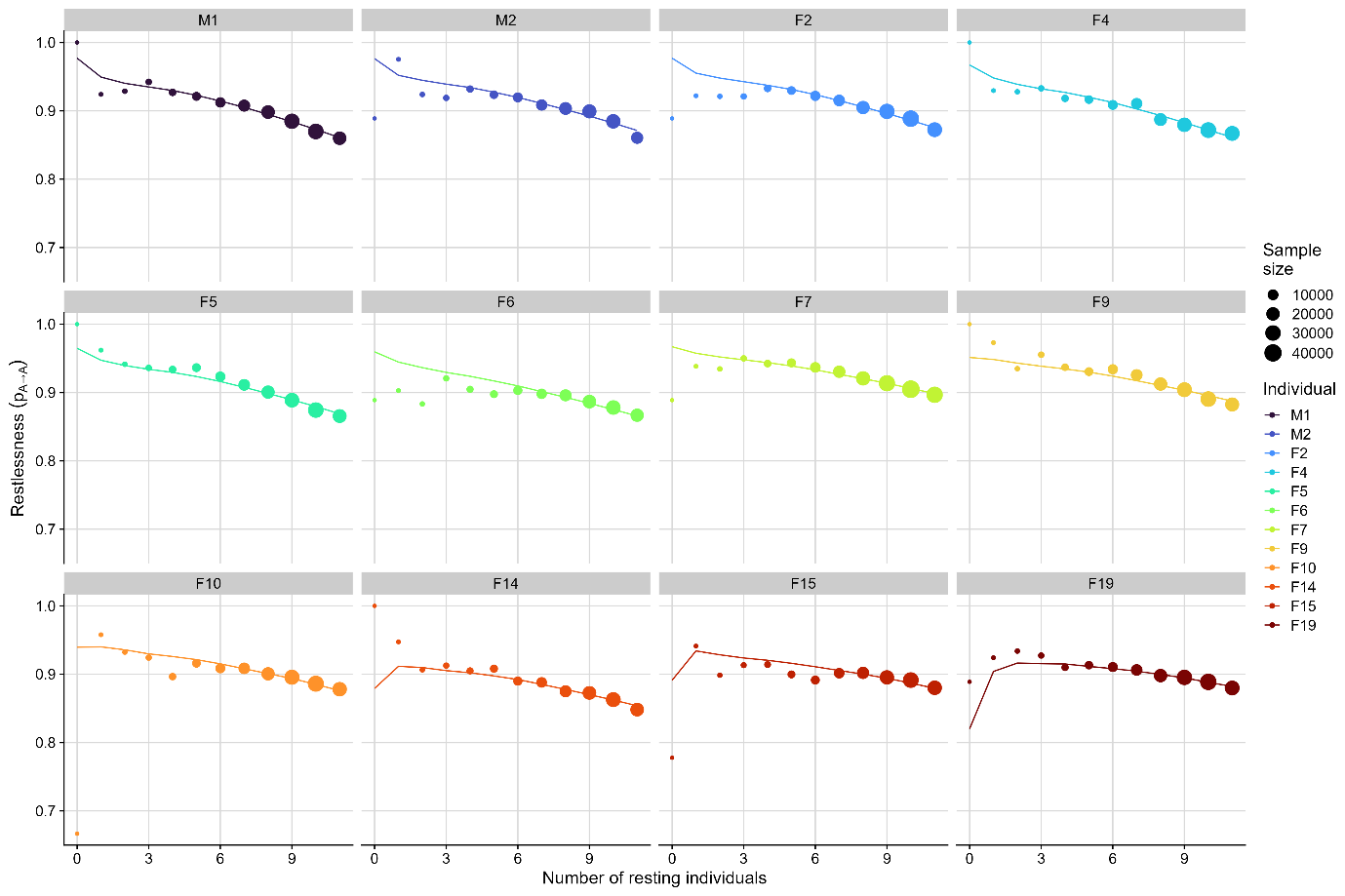

Figure S15 Model predicted wakefulness (lines) compared to raw data (points, size is sample size) for the number of individuals simultaneously resting when conditioning on the focus individual also resting. Every graph is one individual.
